# Functional Analysis of the Expanded Phosphodiesterase Gene Family in *Toxoplasma gondii* Tachyzoites

**DOI:** 10.1101/2021.09.21.461320

**Authors:** William J. Moss, Caitlyn E. Patterson, Alexander K. Jochmans, Kevin M. Brown

## Abstract

*Toxoplasma* motility is both activated and suppressed by 3’-5’ cyclic nucleotide signaling. Cyclic GMP (cGMP) signaling through TgPKG activates motility, whereas cyclic AMP (cAMP) signaling through TgPKAc1 inhibits motility. Despite their importance, it remains unclear how cGMP and cAMP levels are maintained in *Toxoplasma*. Phosphodiesterases (PDEs) are known to inactivate cyclic nucleotides and are highly expanded in the *Toxoplasma* genome. Here we analyzed the expression and function of the 18-member TgPDE family in tachyzoites, the virulent life-stage of *Toxoplasma*. We detected the expression of 11 of 18 TgPDEs, confirming prior expression studies. A knockdown screen of the TgPDE family revealed four TgPDEs that contribute to lytic *Toxoplasma* growth (TgPDE1, TgPDE2, TgPDE5, and TgPDE9). Depletion of TgPDE1 or TgPDE2 caused severe growth defects, prompting further investigation. While TgPDE1 was important for extracellular motility, TgPDE2 was important for host cell invasion, parasite replication, host cell egress, and extracellular motility. TgPDE1 displayed a plasma membrane/cytomembranous distribution, whereas TgPDE2 displayed an endoplasmic reticulum/cytomembranous distribution. Biochemical analysis of TgPDE1 and TgPDE2 purified from *Toxoplasma* lysates revealed that TgPDE1 hydrolyzes both cGMP and cAMP, whereas TgPDE2 was cAMP-specific. Interactome studies of TgPDE1 and TgPDE2 indicated that they do not physically interact with each other or other TgPDEs but may be regulated by kinases and proteases. Our studies have identified TgPDE1 and TgPDE2 as central regulators of tachyzoite cyclic nucleotide levels and enable future studies aimed at determining how these enzymes are regulated and cooperate to control *Toxoplasma* motility and growth.

**Importance:** Apicomplexan parasites require motility to actively infect host cells and cause disease. Cyclic nucleotide signaling governs apicomplexan motility, but it is unclear how cyclic nucleotide levels are maintained in these parasites. In search of novel regulators of cyclic nucleotides in the model apicomplexan *Toxoplasma*, we identified and characterized two catalytically active phosphodiesterases, TgPDE1 and TgPDE2, that are important for *Toxoplasma’*s virulent tachyzoite lifecycle. Enzymes that generate, sense, or degrade cyclic nucleotides make attractive targets for therapies aimed at paralyzing and killing apicomplexan parasites.

## Introduction

Apicomplexan parasites are obligate-intracellular protozoan parasites that cause a variety of deadly diseases including malaria, cryptosporidiosis, and toxoplasmosis. Several species of *Plasmodium* cause human malaria, such as *P. falciparum* and *P. vivax*, resulting in hundreds of thousands of deaths annually (1). *Cryptosporidium parvum and C. hominis* are the main agents of human cryptosporidiosis, a diarrheal disease that kills tens of thousands of children and infants each year (2). *Toxoplasma gondii* (*Toxoplasma*) infections are less deadly, but are much more widespread.

*Toxoplasma* infects and persists within 25-30% of the global human population and causes toxoplasmosis, which can be fatal for immunosuppressed individuals or developing fetuses (3). Although apicomplexan parasites cause distinct diseases in numerous organ systems, lytic parasite growth is the primary source of pathogenesis caused by apicomplexan parasites (4). Therefore, a better understanding of how these parasites progress through their lytic cycles will reveal molecular targets for novel therapeutic interventions.

The apicomplexan asexual lytic cycle occurs in five general steps: attachment to a host cell, invasion, formation of the parasitophorous vacuole (PV), intracellular replication, and egress (5). Attachment comprises interactions between parasite and host surface proteins/glycoproteins. Upon firm attachment, secretion of specialized secretory organelles called micronemes and rhoptries embed a ring-like invasion complex into the host cell plasma membrane that forms a transient portal for parasite entry (6). The parasite glideosome, a surface adhesin-linked actin-myosin motor, provides the locomotive force for invasion and other motile processes (5). During invasion, the PV is formed from host plasma membrane (stripped of host proteins) and provides an interface between the parasite and host for parasite effector export, immune subversion, and nutrient acquisition (7–9). Once inside the PV, parasites undergo asexual replication generating up to several dozens of newly formed parasites (4). To conclude the lytic cycle, parasites secrete pore-forming microneme proteins and upregulate motility to egress from the PV and host cell (10). The parasite lytic cycle directly results in tissue destruction, increased parasite burden, and subsequent inflammation that is the cornerstone of apicomplexan virulence and pathogenesis. Therefore, understanding how parasites modulate motility for lytic growth is critical to understanding how they cause disease.

Apicomplexans have adapted second messenger signaling systems to regulate microneme secretion and motility (11–15). Purine cyclic nucleotides (cGMP and cAMP) and ionic calcium (Ca^2+^) signaling pathways are central regulators of apicomplexan motility, but there are species-dependent variations in how they operate. In general, external signals will stimulate parasite guanylate cyclase(s) to produce cGMP from GTP (16–26). Accumulation of cGMP in the parasite cytosol will activate protein kinase G (PKG), the only known cGMP effector in apicomplexans (27). Catalytically active PKG is required for microneme secretion and motility by stimulating Ca^2+^ flux, and through an unknown mechanism that cannot be bypassed by exogenous Ca^2+^ (28–34). PKG is thought to stimulate release of Ca^2+^ stores by upregulating inositol 1,4,5-trisphosphate (IP3) signaling and by phosphorylating ICM1, a multipass membrane protein essential for PKG-dependent calcium mobilization (35). Cytosolic Ca^2+^ activates Ca^2+^-binding proteins that regulate microneme secretion (e.g. calcium-dependent protein kinases, vesicle fusion machinery) and motility (e.g. calmodulins) (36–38). The role of cAMP signaling is less conserved in Apicomplexa, where cAMP signaling through protein kinase A catalytic subunit (PKAc) acts in concert with cGMP and Ca^2+^ for *Plasmodium* invasion (13, 39–43), yet PKAc1 inhibits motility following invasion by negatively regulating Ca^2+^ in *Toxoplasma* (44, 45). In either scenario, it is clear that cyclic nucleotide levels must be tightly controlled for timely motility in Apicomplexa.

There is growing evidence for the importance of cyclic nucleotide turnover in apicomplexan parasites. Phosphodiesterases (PDEs) inactivate cyclic nucleotides through hydrolysis (46) and are conserved in Apicomplexa (47). Studies using mammalian PDE inhibitors provided the first experimental evidence for the importance of apicomplexan PDEs. Zaprinast and BIPPO upregulate parasite cGMP-dependent motility, while prolonged treatment is lethal (48). Similarly, 3-isobutyl-1-methylxanthine (IBMX) upregulates parasite cAMP and modulates parasite motility, growth and development but its effects are likely species- and life-stage-dependent (47, 49, 50). At the single gene level, most of what is known about apicomplexan PDEs comes from *Plasmodium* research. *Plasmodium* encodes four PDEs (α/β/γ/δ) that all degrade cGMP, with PDEβ (essential in asexual blood stages) also possessing cAMP hydrolytic activity (13, 51). *Cryptosporidium* also encodes a limited set of PDEs (three) but have yet to be characterized. In contrast, *Toxoplasma* encodes 18 PDEs of unknown functional significance. Recent expression studies indicated that the *Toxoplasma* PDE family consists of life-stage dependent PDEs with diverse subcellular localizations (44, 52, 53). Similar to *Plasmodium* PDEβ (42), *Toxoplasma* PDE8 and PDE9 are dual-specific PDEs, although PDE9 was deemed dispensable for tachyzoite growth (53). It is currently unclear which PDEs are primarily responsible for cGMP and/or cAMP turnover in *Toxoplasma* or whether they are functionally redundant.

To facilitate the functional analysis of the *Toxoplasma* PDE family, we created conditional knockdown lines for each TgPDE using a mini-auxin-inducible degron (mAID) system (33). Using this system, we assessed the expression and localization of each TgPDE in the virulent tachyzoite life-stage and measured their contributions to lytic growth following conditional knockdown. We determined that TgPDE1 and TgPDE2 were critical for lytic parasite growth and possessed distinct cyclic nucleotide preferences and subcellular distributions. Furthermore, interactome studies of TgPDE1 and TgPDE2 indicated that they may function in unique signaling complexes and receive novel modes of regulation. Taken together, our studies have identified TgPDE1 and TgPDE2 as central regulators of tachyzoite cyclic nucleotide levels and enable future studies aimed at determining how these enzymes are regulated and cooperate to control motility for lytic growth.

## Results

### *Toxoplasma* encodes 18 putative PDEs with diverse domain architectures

There are 18 phosphodiesterases visible in the *Toxoplasma* genome, but their roles in cyclic nucleotide turnover and parasite fitness are largely undetermined. One or more TgPDEs are suspected to be vital to the tachyzoite lytic lifecycle as treatment with human PDE inhibitors, like zaprinast, blocks plaque formation (48) (Fig. S1).

Conversely, we noted that the non-selective broad-spectrum human PDE inhibitor IBMX does not inhibit tachyzoite growth at high concentration (0.5 mM), indicating that inhibition of host PDEs does not significantly impact parasite growth and that zaprinast likely targets an essential TgPDE or subset of TgPDEs (Fig. S1). Furthermore, a genome-wide CRISPR knockout screen indicated that five TgPDEs are potentially important for tachyzoite fitness *in vitro* (phenotype scores <-1) (54). Similarly, *TgPDE1* and *TgPDE2* are refractory to deletion in tachyzoites providing strong indirect evidence for their importance (44). Collectively, these findings compelled us to investigate the function of the TgPDE family in *Toxoplasma*.

Adhering to the TgPDE nomenclature designated by Vo et al. 2020 (53), we analyzed each TgPDE protein sequence for conserved domains using the NCBI Conserved Domain database (55). We determined that TgPDE proteins ranged from 311 aa (TgPDE14) to 3476 aa (TgPDE18) in length with variable domain architectures. Each TgPDE contained a single C-terminal PDEase_I domain (pfam00233), which hydrolyze 3’5’-cyclic nucleotides such as cAMP and cGMP (Fig. 1). TgPDE4 and TgPDE16 contained portions of an ion-transport domain (pfam00520) suggesting they may also regulate, or be regulated by, ion-transport (Fig. 1). TgPDE2 contained a GAF domain (pfam01590) known to bind cyclic nucleotides and modulate PDE activity (56) (Fig. 1). Low confidence non-specific domain fragments (not shown) were also detected for TgPDE7 (transcriptional regulator ICP4, cl33723), TgPDE9 (transcriptional termination factor Rho, cl36163), TgPDE15 (OCRE, cl23757), and TgPDE18 (MAEBL, cl31754; transcriptional termination factor Rho, cl36163; U2 snRNP auxiliary factor, cl36941). Transmembrane domains within TgPDE proteins were predicted using TOPCONS (57). Except for TgPDE2, TgPDE3, and TgPDE14, all other TgPDEs had 2- 7 transmembrane domains (Fig. 1), suggesting that most TgPDEs are integral membrane proteins. Therefore, the expanded TgPDE family appears well-suited to degrade cyclic nucleotides in a variety of subcellular compartments with diverse modes of regulation and secondary functions.

**Fig 1.**
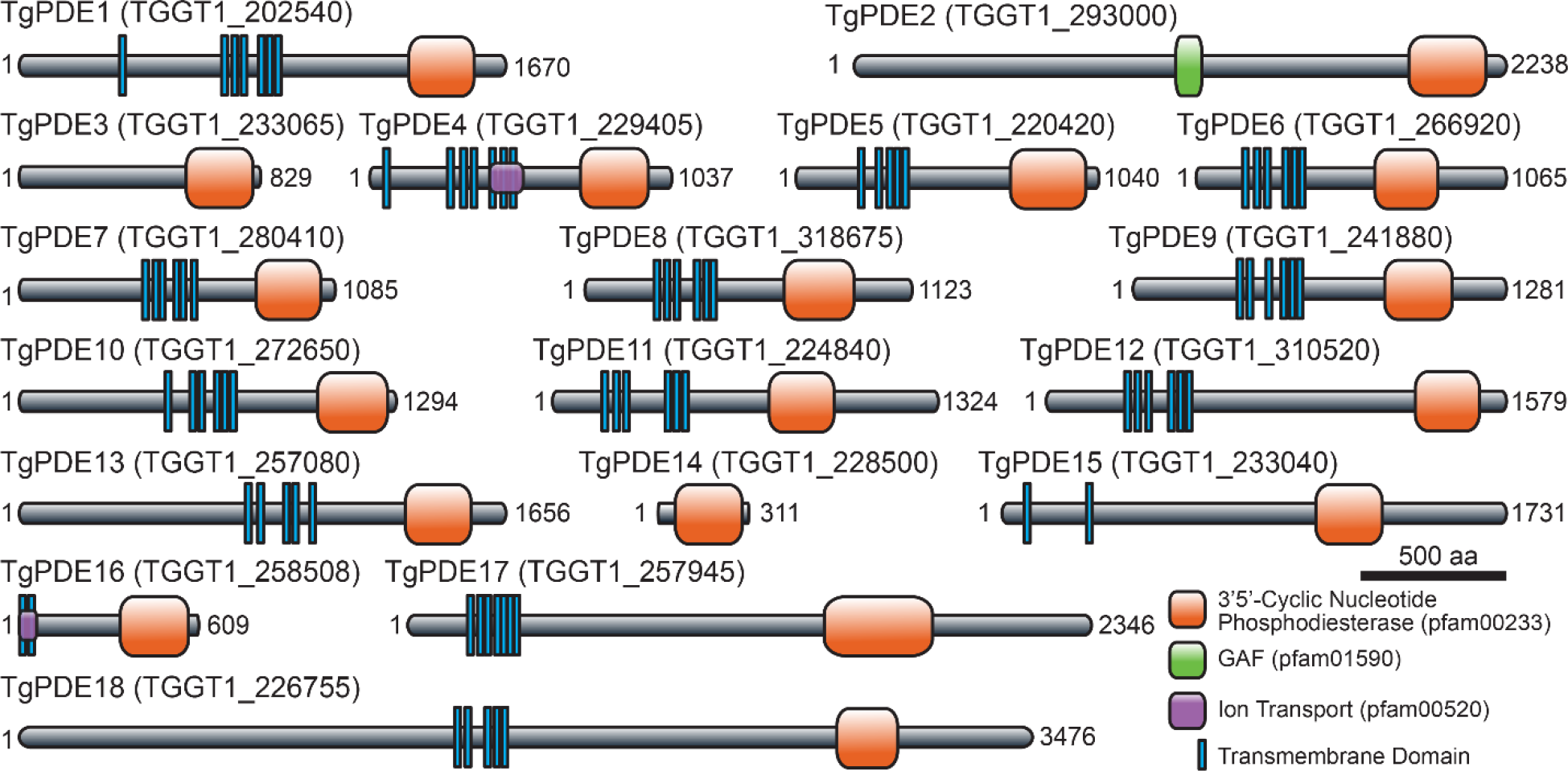
Predicted protein domain architecture of the TgPDE family. Transmembrane domains and conserved protein domains of the 18 putative 3’5’-cyclic nucleotide phosphodiesterases in *T. gondii* based on TOPCONS and NCBI Conserved Domain Database searches.

### Expression and distribution of TgPDEs in tachyzoites

To investigate the expression of each TgPDE in tachyzoites, we implemented an auxin-inducible degron system for detection and conditional depletion of mAID-3HA tagged proteins (33). Starting with RH TIR1-3FLAG, which expresses the plant auxin receptor TIR1, we used CRISPR/Cas9 genome editing (58) to tag each *TgPDE* gene with a short homology-flanked *mAID-3HA, HXGPRT* cassette (Fig. 2A) as described (59). Following drug selection and isolation of clones, genomic DNA was harvested for diagnostic PCRs to distinguish untagged clones from mAID-3HA tagged clones. Our primer design allowed integration validation such that PCR1 should only show a 283- 335 bp band in untagged wildtype genomic DNA, while PCR2 should only show a 493- 534 bp band in TgPDE-mAID-3HA genomic DNA depending on each *TgPDE* gene.

**Fig 2.**
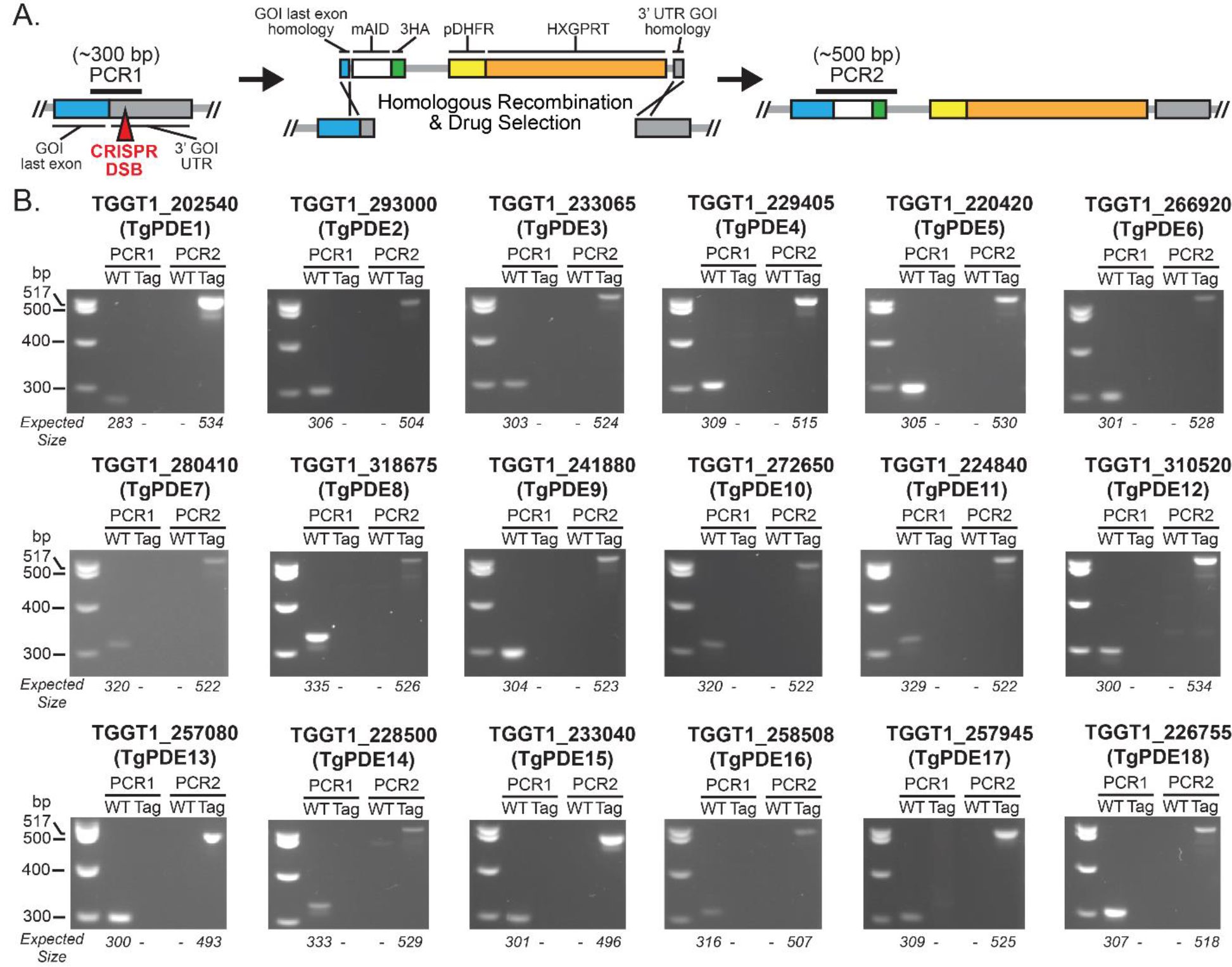
Creation of conditional knockdown lines for all 18 TgPDEs. (A) CRISPR/Cas9 genome editing strategy used to append an *mAID-3HA, HXGPRT* cassette (with 40 bp homology flanks) to the 3’ end of each *TgPDE* gene in the RH TIR1-3FLAG background. (B) Diagnostic PCR confirmation of successful *mAID-3HA, HXGPRT* integration for each *TgPDE* gene. Differential genomic positions of PCR1 and PCR2 are shown in (A). WT, wild type parental line RH TIR1-3FLAG. Tag, specific RH TgPDE- mAID-3HA clone.

Diagnostic PCR revealed that each *TgPDE* was successfully tagged with the *mAID- 3HA, HXGPRT* for detection and functional analyses (Fig. 2B).

To determine which TgPDEs are expressed in tachyzoites, we used immunofluorescence (IF) microscopy and immunoblotting to detect the 3HA epitope for each TgPDE-mAID-3HA fusion. TgPDE1, 2, 5, 6, 7, 9, 10, 11, 12, 13, and 18 showed expression in tachyzoites by IF microscopy and/or immunoblotting (Fig. 3). We were unable to detect TgPDE8 by either method, which can be detected with the more sensitive spaghetti monster-HA (smHA) tag in tachyzoites (53). Of the TgPDEs detected by IF microscopy, we observed diverse subcellular distribution patterns reminiscent of plasma membrane (TgPDE1, 7, 9, 10), endoplasmic reticulum (TgPDE2, 11, 13, 18), mitochondrion (TgPDE5), nucleus (TgPDE6), apical cap (TgPDE9), and cytomembranes (TgPDE1, 2, 6, 10, 11, 13, 18) (Fig. 3A). Co-staining with markers for each compartment and super-resolution microscopy will be needed to precisely localize each TgPDE going forward. Of the TgPDEs detected by immunoblotting, all could be detected in full-length, but with smaller isoforms also detected for TgPDE2, 7, and 10 indicating that they could be regulated through alternative splicing or proteolysis (Fig. 3B). To determine whether auxin-treatment could deplete the TgPDE-mAID-3HA fusions, parasites were treated with vehicle or 0.5 mM auxin (3-indoleacetic acid; IAA) for 18 h, lysed, and analyzed by SDS-PAGE with immunoblotting for the 3HA epitope. We observed that all detectable TgPDE-mAID-3HA fusions were efficiently depleted following auxin treatment (Fig. 3B). Taken together, we have identified at least 11 TgPDEs that may regulate cyclic nucleotide turnover in tachyzoites and have developed a robust knockdown system for assessing the function of each member of the TgPDE family.

**Fig 3.**
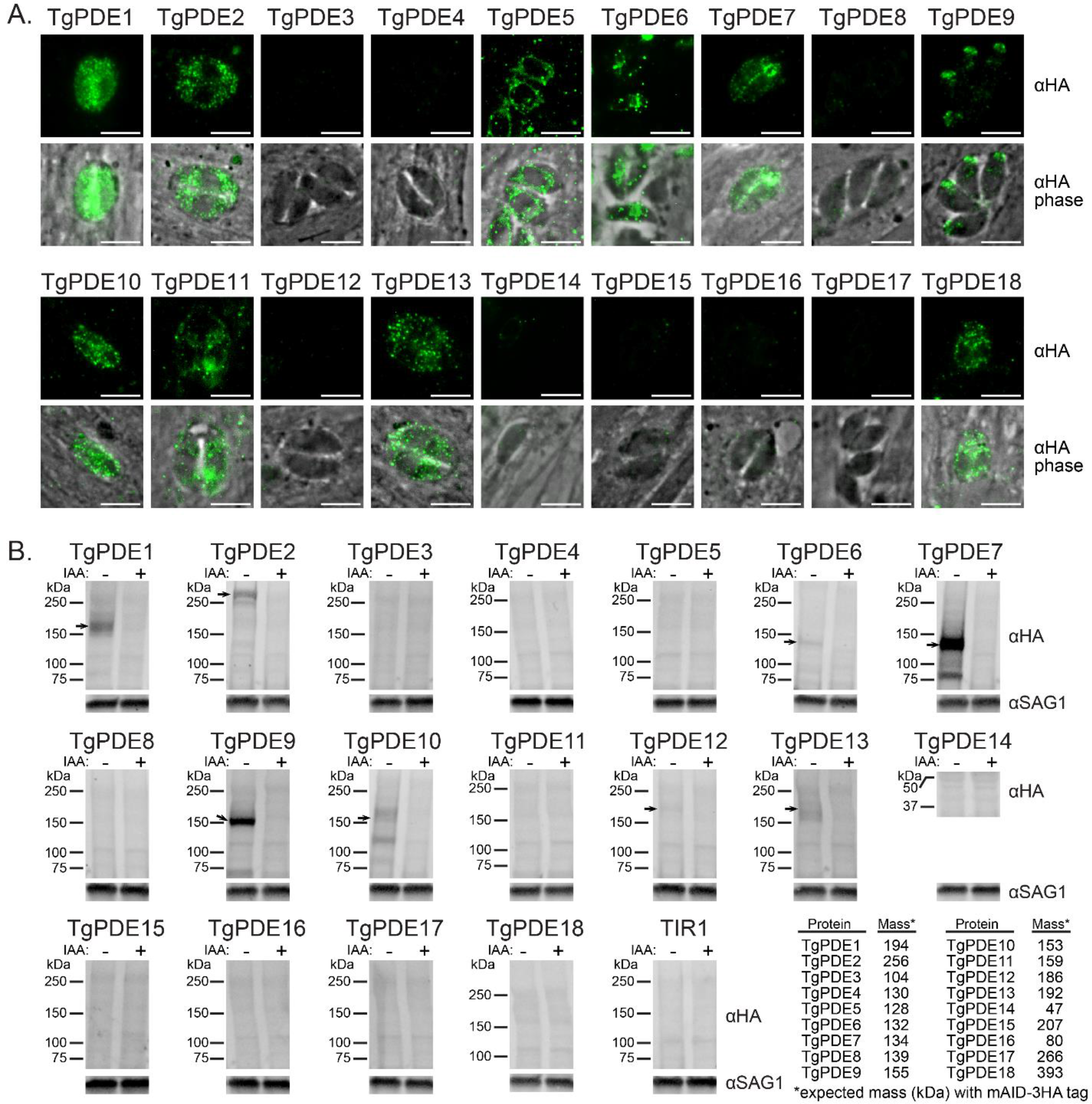
Expression and depletion of TgPDE-mAID-3HA fusions in tachyzoites. (A) IF microscopy of intracellular RH TgPDE-mAID-3HA parasites labeled with mouse anti-HA and goat anti-mouse IgG Alexa Fluor 488. Scale bar = 5 μm. (B) Immunoblots of lysates from RH TIR1-3FLAG (TIR1) and RH TgPDE-mAID-3HA (TgPDE) parasites treated with vehicle (EtOH) or 0.5 mM IAA for 18 h. Blots probed with mouse anti-HA, rabbit anti-SAG1, goat anti-mouse-AFP800, and goat anti-rabbit-AFP680. Table indicates predicted total mass of each TgPDE including their mAID-3HA tag (12 kDa). Arrows indicate IB-detectable TgPDE-mAID-3HA fusions in tachyzoites.

### Four TgPDEs contribute to the tachyzoite lytic lifecycle

To assess the individual contribution of each TgPDE to overall tachyzoite fitness, we performed plaque assays for all 18 TgPDE conditional knockdown mutants. We reasoned that even TgPDEs expressed below the limit of detection may still serve important functions, so they were also included in the screen. Freshly-harvested RH TIR1-FLAG and RH PDE-mAID-3HA tachyzoites were inoculated onto HFF monolayers, treated with 0.5 mM IAA (to deplete mAID-3HA fusions) or vehicle for 8 days, fixed with ethanol, and stained with crystal violet to visualize plaque formation (e.g. macroscopic zones of clearance on host cell monolayers). Since plaque formation requires the completion of several rounds of lytic growth, defects in any step of the lytic cycle will reduce plaque formation. As previously reported (33, 38), IAA did not affect the ability of RH TIR1-3FLAG to form plaques (Fig. 4). However, we observed that knockdown of four TgPDEs caused significant plaquing defects compared to vehicle treatment (Fig. 4). Depletion of TgPDE5 and TgPDE9 reduced plaque area by 32% and 26%, respectively, indicating suboptimal lytic growth (Fig. 4). In contrast, depletion of TgPDE1 and TgPDE2 significantly reduced both plaque area and the number of plaques formed (Fig. 4). TgPDE1 depletion reduced plaque area by 58%, indicating that is critical for tachyzoite growth. TgPDE2 depletion reduced plaque area by 95%, indicating that it may be a master regulator of cAMP and/or cGMP turnover in tachyzoites. The non-redundant phenotypes associated with separate TgPDE1 and TgPDE2 depletion suggests that they may have unique cyclic nucleotide preferences which would lead to opposing roles in regulating motility for lytic cycle progression.

**Fig 4.**
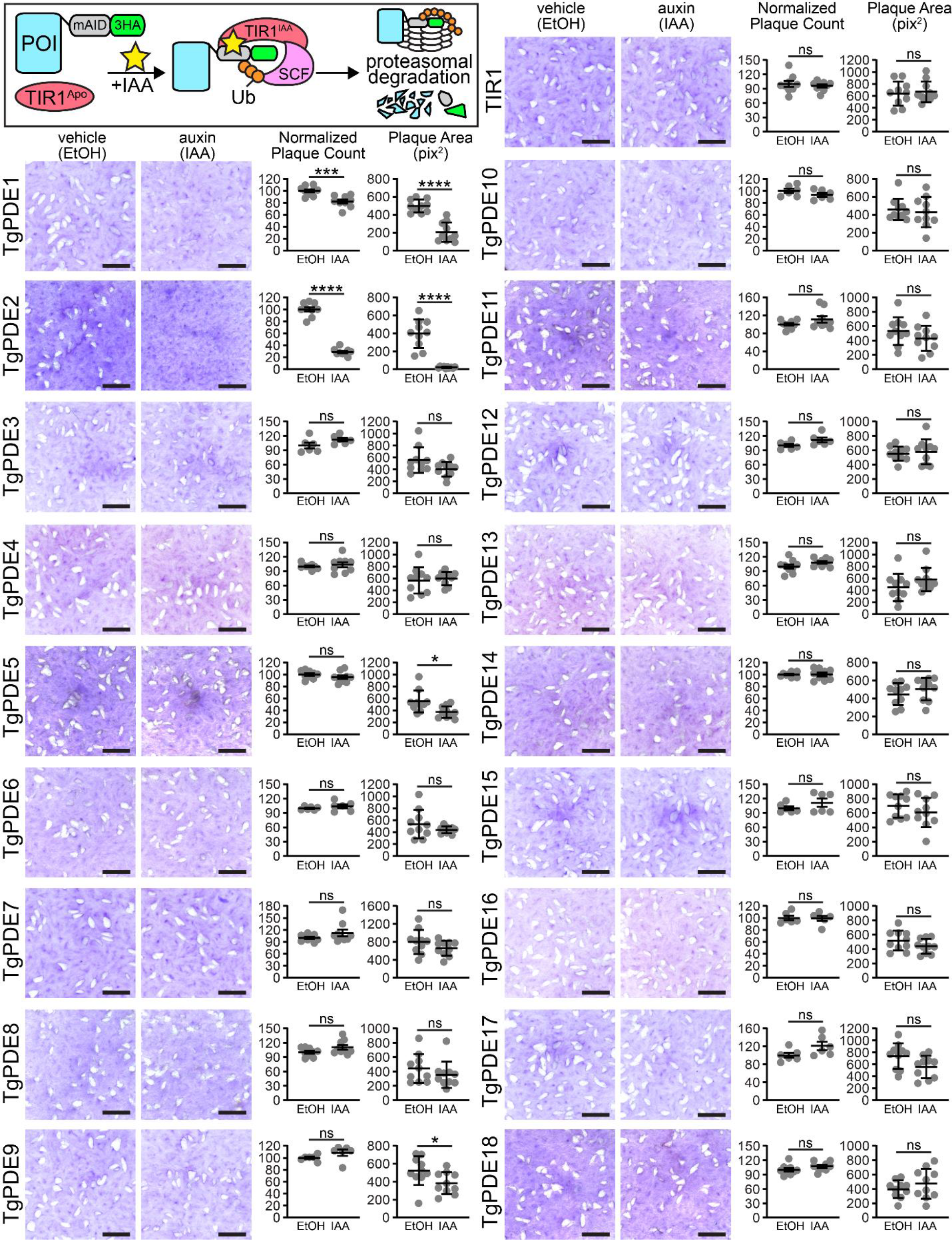
Contribution of each TgPDE to tachyzoite growth. Inset: Auxin-inducible protein degradation. Plaques formed on HFF monolayers by RH TIR1-3FLAG (TIR1) or derivative RH TgPDE-mAID-3HA lines treated with vehicle (EtOH) or 0.5 mM IAA for 8 days. Scale bar = 5 mm. Plaque count data represented as mean ± SD (n = 6 or 9) replicates combined from N = 2 or 3 trials, respectively. For baseline consistency, plaque counts were normalized to 100 vehicle treated plaques for each line. Plaque area data represented as mean ± SD (n = 10) from the representative images shown. Unpaired student’s t test (ethanol vs IAA), *P*: * ≤ 0.05, *** ≤ 0.001, **** ≤ 0.0001.

### Biochemical activities of TgPDEs

To determine the substrate preference of each TgPDE, we first generated recombinant 6HIS-SUMO fusions of TgPDE catalytic domain fragments (6HIS-SUMO- TgPDE^CAT^) from *E. coli*. This strategy was chosen to facilitate expression, solubility, and purification of non-denatured TgPDE^CAT^ fragments based on the retained activity of 5HIS-PfPDEα^CAT^, an analogous fragment of a cGMP-specific PDE from *P. falciparum* purified from *E. coli* (47). For *E. coli* expression, TgPDE^CAT^ fragments (Fig. S2A) were cloned from GT1 tachyzoite cDNA libraries or synthesized as dsDNA gene fragments and ligated into pET*-6HIS-SUMO* (60). Similar constructs containing PfPDEα^CAT^ and PfPDEβ^CAT^ or HsGSDMD (human gasdermin D) served as positive and negative controls for PDE activity, respectively. The sequence-verified expression constructs were then transformed into SHuffle T7 *E. coli*, induced with isopropylthio-β-galactoside (IPTG), and 6HIS-SUMO protein fusions were purified from sarkosyl-solubilized inclusion bodies using immobilized metal affinity chromatography with Ni-NTA gravity flow columns (Fig. S2B). The recombinant protein fractions were analyzed by SDS-PAGE with total protein staining and we determined that each 6HIS-SUMO-TgPDE^CAT^ protein was captured and adequately purified for downstream analysis (Fig. S3). To determine whether the recombinant 6HIS-SUMO-TgPDE^CAT^ proteins possessed PDE activity, we performed PDE-Glo assays using cAMP or cGMP as the relevant substrate (Fig. S4A). Unfortunately, we were unable to reliably detect PDE activity for the 6HIS- SUMO-TgPDE^CAT^ proteins with even 2 h reactions at 37°C (Fig. S4B, S4C). These results suggest that generating catalytically-active recombinant TgPDE^CAT^ proteins will require further optimization (detergents, refolding, etc.) or that TgPDEs need to be purified in full-length from *Toxoplasma* to retain activity.

To determine whether endogenously-expressed TgPDEs are active, we focused on the two most critical TgPDEs for tachyzoite growth: TgPDE1 and TgPDE2 (Fig. 4). We used immunoprecipitation (IP) to capture TgPDE1-mAID-3HA and TgPDE2-mAID- 3HA from native soluble tachyzoite lysates using anti-HA magnetic chromatography with HA peptide elution (Fig. 5A). IP fractions for total protein, soluble protein, and eluted protein were analyzed by immunoblotting and we observed that TgPDE1 and TgPDE2 were effectively captured (Fig. 5B). We believe the non-denaturing detergents in the native lysis buffer interacted with the transmembrane domains of TgPDE1 and slowed its migration during SDS-PAGE, as has been described for other transmembrane proteins (61, 62). In agreement, the supershifted species of TgPDE1-mAID-3HA was not observed when lysates were prepared without these detergents (Fig. 3B). Due to the nature of the HA peptide elution, and relatively low yield of immunoprecipitated proteins, we were unable to measure the exact concentrations of captured proteins. Therefore, we performed PDE-Glo assays using equivalent volumes of IP elution fractions with the untagged parental line elution fraction serving as the background threshold for PDE activity. Upon addition of 200 nM cAMP substrate, eluted TgPDE1 and TgPDE2 fractions were clearly capable of hydrolyzing cAMP as compared to the negative control and standards (Fig. 5C). When 20 μM cGMP substrate was supplied, only eluted TgPDE1 was able to hydrolyze cGMP (Fig. 5D).

**Fig 5.**
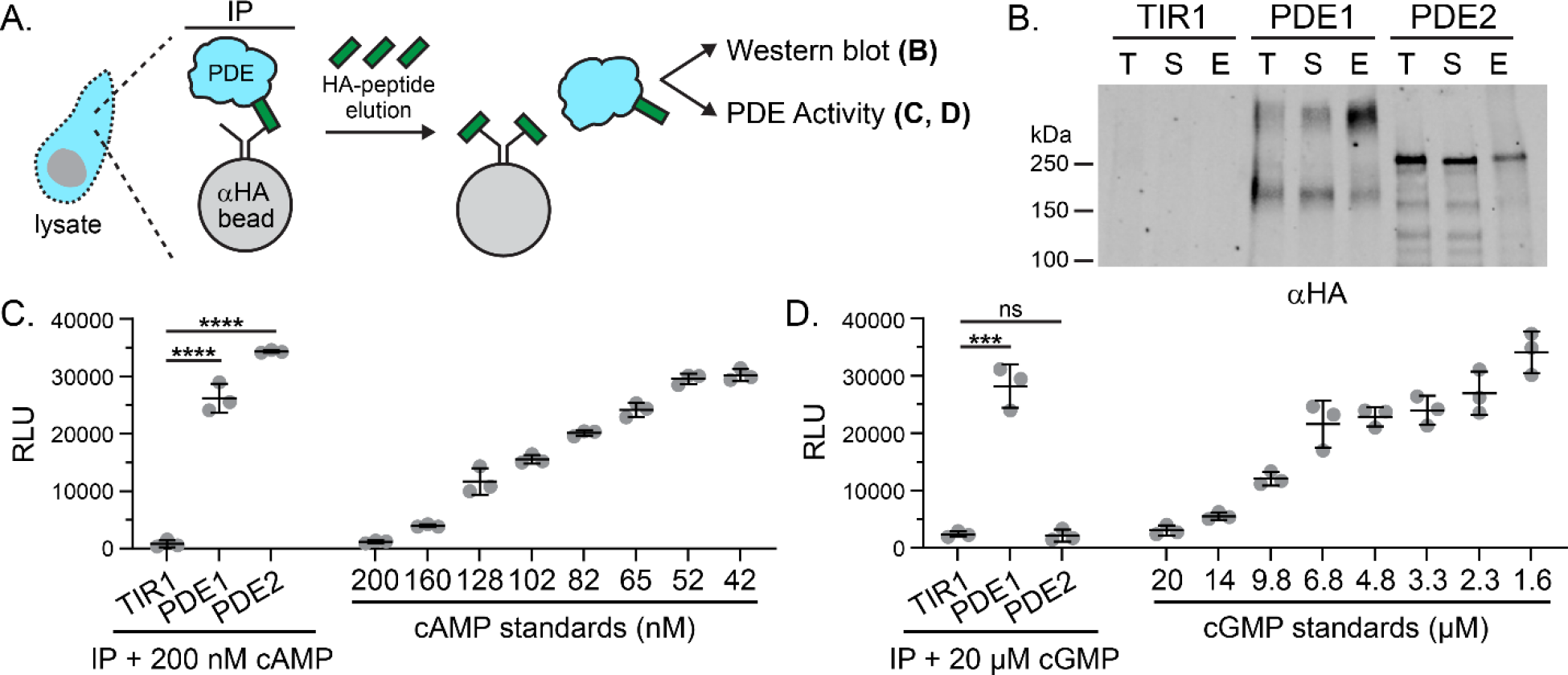
Phosphodiesterase activity of immunoprecipitated TgPDE1 and TgPDE2. (A) Immunoprecipitation strategy for purifying tagged TgPDE1 and TgPDE2 proteins from tachyzoite native lysates. Untagged parental line RH TIR1-3FLAG (TIR1) served as negative control. (B) Representative immunoblot of immunoprecipitation fractions probed with rat anti-HA and goat anti-rat IRDye 800CW. T = total lysate, S = soluble lysate, E = eluate. (C) Cyclic AMP phosphodiesterase activity of immunoprecipitated elution fractions incubated 1:1 with 0.2 µM cAMP for 2 h at 37°C. Standards shown were incubated 1:1 with PDE storage buffer for 2 h at 37°C. (D) Cyclic GMP phosphodiesterase activity of immunoprecipitated elution fractions incubated 1:1 with 20 µM cGMP for 2 h at 37°C. Standards shown were incubated 1:1 with PDE storage buffer for 2 h at 37°C. (C,D) Data represented as mean ± SD (n = 3) from one of four trials (N=4) with similar outcomes. Statistical significance was determined using an unpaired student’s t test (TIR1 vs TgPDE1; TIR1 vs TgPDE2), *P*: *** ≤ 0.001, **** ≤ 0.0001.

To determine if any other TgPDEs were unexpectedly pulled down with TgPDE1- mAID-3HA or TgPDE2-mAID-3HA, we performed similar IP experiments without HA peptide elution and analyzed all captured proteins by liquid chromatography mass spectrometry (LC-MS/MS) (Fig. S5A). In this case, RH YFP-AID-3HA served as a control for non-specific protein interactions. IP fractions were first analyzed by total protein staining and immunoblotting (Fig. S5B,C). While the relative abundance of captured protein was low (Fig. S5B), we were able to clearly detect the target proteins by immunoblotting (Fig. S5C). TgPDE1-mAID-3HA and TgPDE2-mAID-3HA both retained hydrolytic activity while immobilized on anti-HA magnetic beads, indicating that native protein complexes were captured (Fig. S5D). Importantly, no other TgPDE co- immunoprecipitated with either TgPDE1 or TgPDE2 in two independent trials (Fig. S5E, Table S4). Furthermore, formaldehyde-crosslinking protein interactions prior to parasite lysis and IP (XL-LC-MS/MS) did not stabilize interactions between TgPDE1 and TgPDE2 or any other TgPDE (Table S5). However, these interactome studies indicate that TgPDE1 and TgPDE2 are potentially regulated by other proteins through protein-protein interactions and/or post-translational modifications.

### TgPDE1 and TgPDE2 regulate distinct steps in the lytic cycle

Given that TgPDE1 and TgPDE2 appear non-redundant for tachyzoite growth and have distinct substrates, we speculated that they control distinct steps within the lytic cycle. To test this hypothesis, we performed standard assays for tachyzoite replication, host cell attachment and invasion, host cell egress, and extracellular motility.

To determine whether TgPDE1 and TgPDE2 are important for parasite replication, freshly-harvested RH TIR1-3FLAG, RH TgPDE1-mAID-3HA, and RH TgPDE2-mAID-3HA parasites were inoculated onto HFF monolayers for 2 h and then treated with vehicle (EtOH) or 0.5 mM IAA. At 24 h post infection, the cultures were fixed and immunolabeled with antibodies to TgSAG1, TgGRA7 and the DNA dye Hoechst 33258. Using IF microscopy, we observed that depletion of TgPDE2, but not TgPDE1, reduced the size of the parasite vacuoles and increased the number of irregular vacuoles (Fig. 6A), which contain an atypical number of parasites (non 2N) or parasites with abnormal morphology. To quantify the replication defects, we counted the numbers of regular and irregular vacuoles, and the numbers of parasites within each.

**Fig 6.**
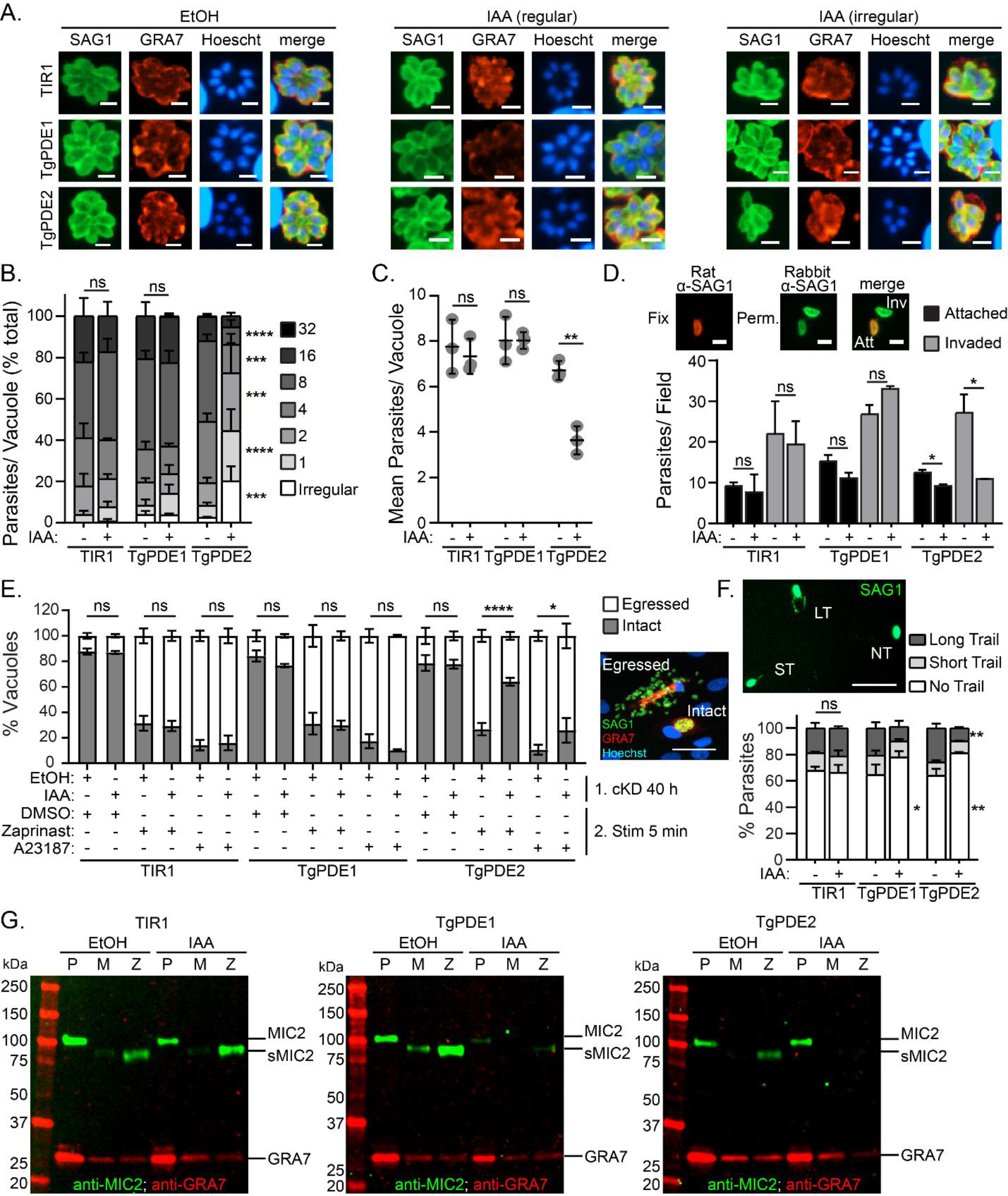
Role of TgPDE1 and TgPDE2 in the tachyzoite lytic cycle. RH TIR1-3FLAG (TIR1) and derivative RH PDE-mAID-3HA parasites (TgPDE1; TgPDE2) were analyzed for defects in the lytic cycle following conditional knockdown of TgPDE1 or TgPDE2 with auxin. (A-C) Replication assay of parasites grown in HFFs treated with vehicle (EtOH) or 0.5 mM IAA for 22 h from N=3 independent trials. (A) IF microscopy of fixed parasites labeled with antibodies to TgSAG1, TgGRA7 and the DNA dye Hoechst 33258. Scale bar = 5 μm. Irregular vacuoles contain an irregular number of parasites or parasites with abnormal morphology. (B) Distribution of parasites per vacuole ± SD. For each line, percentages of parasites in each bin were compared between EtOH and IAA treatment and statistical significance was determined using a two-way ANOVA with Tukey’s multiple comparisons test. *P*: *** ≤ 0.001, **** ≤ 0.0001. (C) Mean parasites per vacuole ± SD. Unpaired student’s t test (EtOH vs IAA), *P*: ** ≤ 0.01. (D) Parasite invasion of HFFs (20 min) following treatment with EtOH or 0.5 mM IAA for 16 h from N=2-3 independent trials. Shown are the mean numbers of attached and invaded parasites per field ± SD as determined by IF microscopy using differential permeabilization and immunolabeling with rat and rabbit TgSAG1 antibodies. Scale bar = 5 μm. Unpaired student’s t test (EtOH vs IAA), *P*: * ≤ 0.05. (E) Parasite egress from HFFs (40 h) following treatment with EtOH of 0.5 mM IAA for 22 h and stimulation with vehicle (DMSO), 0.5 mM zaprinast, or 2 μM A23187 for 5 min from N=3 independent trials. Shown are the mean percentages of intact and egressed vacuoles ± SEM based on IF microscopy using immunolabeling with antibodies against TgSAG1 and TgGRA7. Scale bar = 100 μm. Two-way ANOVA with Tukey’s multiple comparisons test. *P*: * ≤ 0.05, **** ≤ 0.0001. (F) Parasite motility on BSA-coated cover glass in EC buffer following 14 h treatment with EtOH or 0.5 mM IAA. Following a 20 min incubation, parasites were fixed to the cover glass and immunolabeled with TgSAG1 antibodies to detect the parasites and their motility trails by IF microscopy. Shown are mean percentages of parasites ± SD with no trail, a short trail (< 1 body length), or a long trail (> 1 body length) from one of two independent trials. Scale bar = 30 μm. Two-way ANOVA with Tukey’s multiple comparisons test. *P*: * ≤ 0.05, ** ≤ 0.01. (G) Microneme secretion assay following 14 h treatment with EtOH or 0.5 mM IAA and stimulation with DMSO or 0.5 mM zaprinast in EC buffer for 10 min as detected by immunoblotting with TgMIC2 and TgGRA7 (constitutively-secreted control protein) antibodies. Representative immunoblots are shown. P = 10% of unstimulated parasite pellet lysate; M = mock-stimulated secreted fraction; Z = zaprinast-stimulated secreted fraction.

Depletion of TgPDE1 did not alter the frequency of irregular vacuoles (Fig. 6B) or the number of parasites within each vacuole (Fig. 6B, 6C). However, depletion of TgPDE2 significantly increased the frequency of irregular vacuoles (Fig. 6B) and decreased the number of parasites in each vacuole (Fig. 6B, 6C). Since TgPDE2 is capable of degrading cAMP, loss of TgPDE2 could possibly elevate cAMP, resulting in hyperactivation of TgPKAc1. Overexpression of this protein kinase has previously been shown to cause replication defects when overexpressed (44). Since TgPDE1 was dispensable for replication, similar to loss of TgGC (21) and TgPKG (33), it may preferentially regulate cGMP for motility in tachyzoites even though it is capable of degrading both cGMP and cAMP *in vitro*.

To test whether TgPDE1 and TgPDE2 are important for host cell attachment and invasion, we first treated RH TIR1-3FLAG, RH TgPDE1-mAID-3HA, and RH TgPDE2- mAID-3HA cultures with vehicle (EtOH) or 0.5 mM IAA for 16 h. Following knockdown, the treated parasites were purified and allowed to invade fresh HFF monolayers ± IAA. After 20 min, non-attached parasites were removed and the cultures were fixed.

Extracellular and intracellular parasites were differentially labeled with TgSAG1 antibodies and visualized by IF microscopy (Fig. 6D). Surprisingly, TgPDE1 depletion did not significantly affect attachment or invasion. Instead, we found that depletion of TgPDE2 significantly reduced attachment (26% reduction) and invasion (60% reduction) (Fig. 6D). Elevation of cAMP in extracellular tachyzoites by TgPDE2 depletion could lead to premature activation of TgPKAc1, which is known to rapidly suppress *Toxoplasma* motility following invasion (44, 45).

Like invasion, *Toxoplasma* egress requires cGMP and Ca^2+^ signaling. These signals are naturally activated following parasite replication but can also be stimulated by agonists such as the PDE inhibitor zaprinast (cGMP agonist that also elevates Ca^2+^) and A23187 (Ca^2+^ agonist). Conversely, cAMP signaling is thought to antagonize egress because genetic inhibition of TgPKAc1 causes premature egress (44, 45). To determine whether TgPDE1 and TgPDE2 are required for egress, we performed an assay that measures natural- and agonist-stimulated egress (21). RH TIR1-3FLAG, RH TgPDE1-mAID-3HA, and RH TgPDE2-mAID-3HA were inoculated onto HFFs for 18 h, then treated with vehicle (EtOH) or 0.5 mM IAA to deplete mAID-3HA-tagged proteins. At 40 h, parasites were stimulated with a short pulse of vehicle (DMSO), 0.5 mM zaprinast, or 2 µM A23187. Next, the parasite cultures were fixed and immunolabeled with TgSAG1 and TgGRA7 antibodies to detect parasites and vacuoles by IF microscopy (Fig. 6E). At this timepoint, between 12-21% of parasite vacuoles had egressed naturally. By contrast, treatment with zaprinast or A23187 stimulated parasite egress up to 73% and 86%, respectively. Depletion of TgPDE1 increased the percentage of naturally egressed vacuoles from 16% to 23%, but this change was not statistically significant. Modest increases were also observed for stimulated egress when TgPDE1 was depleted, but the differences were also not statistically significant. TgPDE2 depletion did not affect natural egress at 40 h. However, TgPDE2 depletion significantly antagonized zaprinast- and A23187-stimulated egress. Depletion of TgPDE2 reduced zaprinast-induced egress from 73% to 35% while A23187-induced egress was only reduced from 89% to 74%. These results suggest that TgPDE2 promotes egress by antagonizing cAMP activation of TgPKAc1. In support, inhibition of cGMP signaling through TgPKG blocks premature egress induced by TgPKAc1 genetic inhibition (44).

Following egress, tachyzoites migrate to new host cells using gliding motility (5). To determine whether TgPDE1 and TgPDE2 regulate motility, we first treated RH TIR1- 3FLAG, RH TgPDE1-mAID-3HA, and RH TgPDE2-mAID-3HA cultures with vehicle (EtOH) or 0.5 mM IAA for 14 h. Following knockdown, the treated parasites were purified and resuspended in extracellular (EC) buffer ± EtOH or 0.5 mM IAA. Parasites were allowed to migrate on BSA-coated wells for 20 min, then fixed and immunolabeled with TgSAG1 antibodies to detect the parasites and their motility trails by IF microscopy. With vehicle treatment, 32-36% of parasites displayed motility trails, which were categorized as short (< 1 body length) or long (> 1 body length) (Fig. 6F). Depletion of TgPDE1 decreased the percentage of motile parasites from 36% to 23% (Fig. 6F).

Depletion of TgPDE2 decreased the percentage of motile parasites from 36% to 19%, and the percentage of parasites with long trails from 26% to 10% (Fig. 6F). With respect to their substrate specificities, these results suggest that TgPDE1 and TgPDE2 cooperate to fine-tune cyclic nucleotide levels for proper motility.

Extracellular migration of *Toxoplasma* requires secretion of adhesins (e.g. TgMIC2) from microneme vesicles, permitting substrate-based motility (5). Microneme secretion is dependent on cGMP and Ca^2+^ signaling, which can be strongly upregulated by zaprinast (cGMP agonist) treatment. To better understand why depletion of TgPDE1 and TgPDE2 reduces *Toxoplasma* motility, we examined microneme secretion following TgPDE1 and TgPDE2 knockdown. RH TIR1-3FLAG, RH TgPDE1-mAID-3HA, and RH TgPDE2-mAID-3HA cultures were treated with vehicle (EtOH) or 0.5 mM IAA for 14 h.

Following knockdown, the treated parasites were purified and resuspended in EC buffer ± EtOH or 0.5 mM IAA and incubated with vehicle (DMSO) or 0.5 mM zaprinast for 10 min to facilitate microneme secretion. Parasite pellets and secreted fractions were collected and analyzed by immunoblotting for TgMIC2 and TgGRA7, a constitutively secreted dense granule protein. In vehicle-treated parasites, zaprinast upregulated TgMIC2 secretion compared to mock-stimulation (DMSO) (Fig. 6G). Surprisingly, depletion of TgPDE1 reduced the amount of TgMIC2 available for secretion, but not the capacity to secrete this smaller pool of TgMIC2 by zaprinast treatment (Fig. 6G). A reduction of microneme proteins within the parasite may be due to premature secretion if cGMP is elevated by TgPDE1 depletion or a general fitness defect. Depletion of TgPDE2 blocked both basal- and zaprinast-induced microneme secretion (Fig. 6G).

Thus, the motility defects caused by TgPDE1 and TgPDE2 depletion are likely caused by aberrant secretion of micronemes.

Collectively, our investigation of the TgPDE family revealed that TgPDE1 is a dual-specific PDE and TgPDE2 is a cAMP-specific PDE that are critical for tachyzoite lytic growth and motility.

## Discussion

Recent studies have demonstrated that purine 3’-5’ cyclic nucleotides, cGMP and cAMP, are master regulators of lytic lifecycle progression in *Toxoplasma* (12, 15). While prior studies focused on the biosynthesis and effector functions of these signaling molecules, here we focused on an expanded family of PDEs to identify regulators of cyclic nucleotide turnover in *Toxoplasma* (summarized in Table 1). Comparative genomics revealed that coccidian apicomplexans encode 8-19 PDE genes, the most out of the four principle apicomplexan groups (gregarina, cryptosporidia, hematozoa, coccidia) (https://www.veupathdb.org/). *Toxoplasma* serves as a tractable model for investigating coccidian PDEs since it encodes 18 representative PDEs. *Toxoplasma* PDEs have a core domain architecture consisting of a C-terminal PDEase_I domain and variable N-terminal features, although mutational analysis will be needed to identify domains necessary for TgPDE function. By tagging each TgPDE gene with a regulatable epitope tag, we were able to detect 11 TgPDEs expressed in tachyzoites with a variety of subcellular distributions. Our conditional knockdown screen of the TgPDE family revealed that four TgPDEs (TgPDE1, 2, 5, 9) independently contribute to tachyzoite growth. From these, TgPDE1 and TgPDE2 emerged as principle regulators of tachyzoite fitness. TgPDE1 and TgPDE2 displayed distinct subcellular localizations, cyclic nucleotide preferences, and interactomes, suggesting opposing roles in cyclic nucleotide regulation in tachyzoites. Accordingly, investigation of specific steps within the lytic cycle revealed unique phenotypes associated depletion of TgPDE1 or TgPDE2.

**Table 1.**
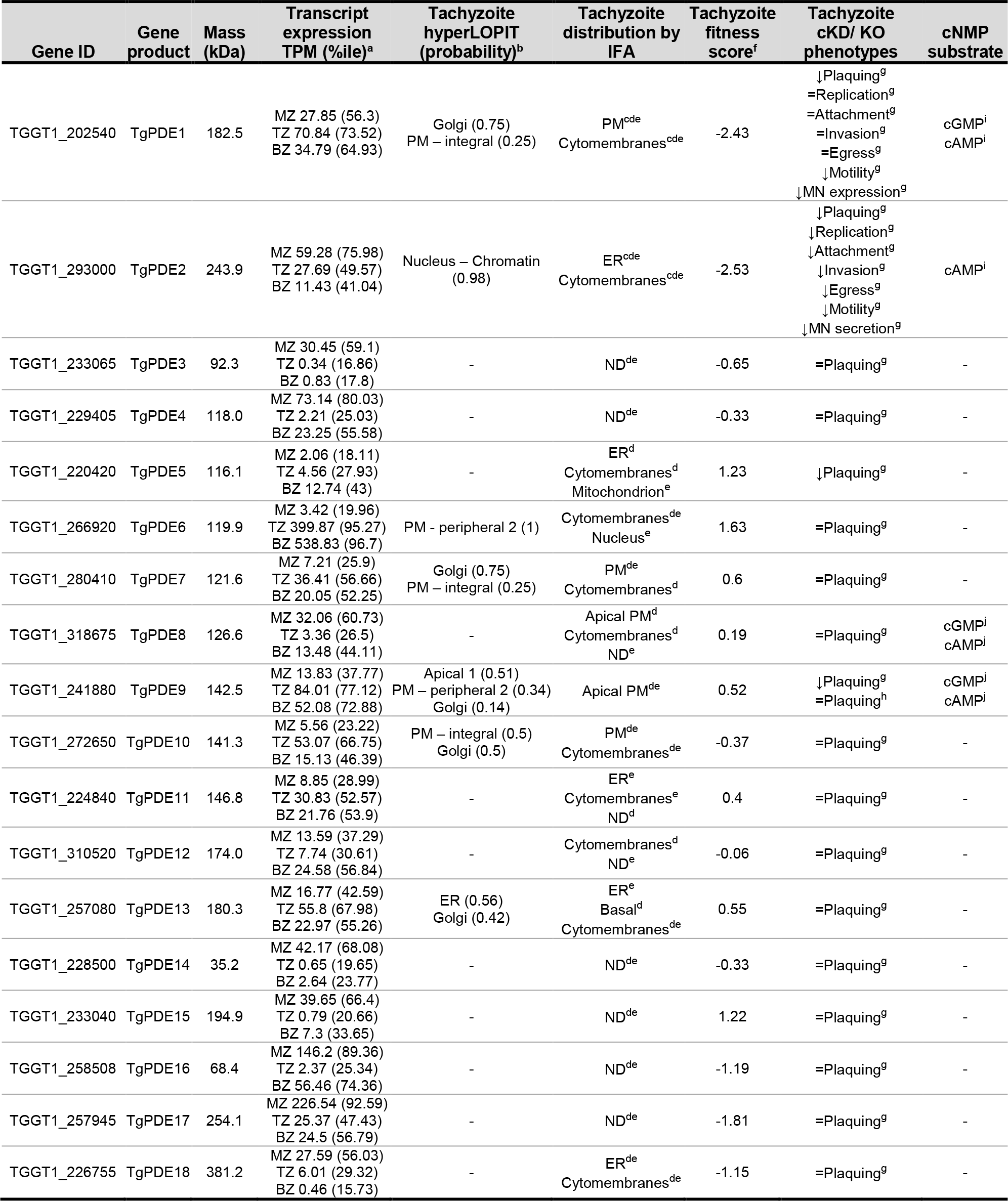
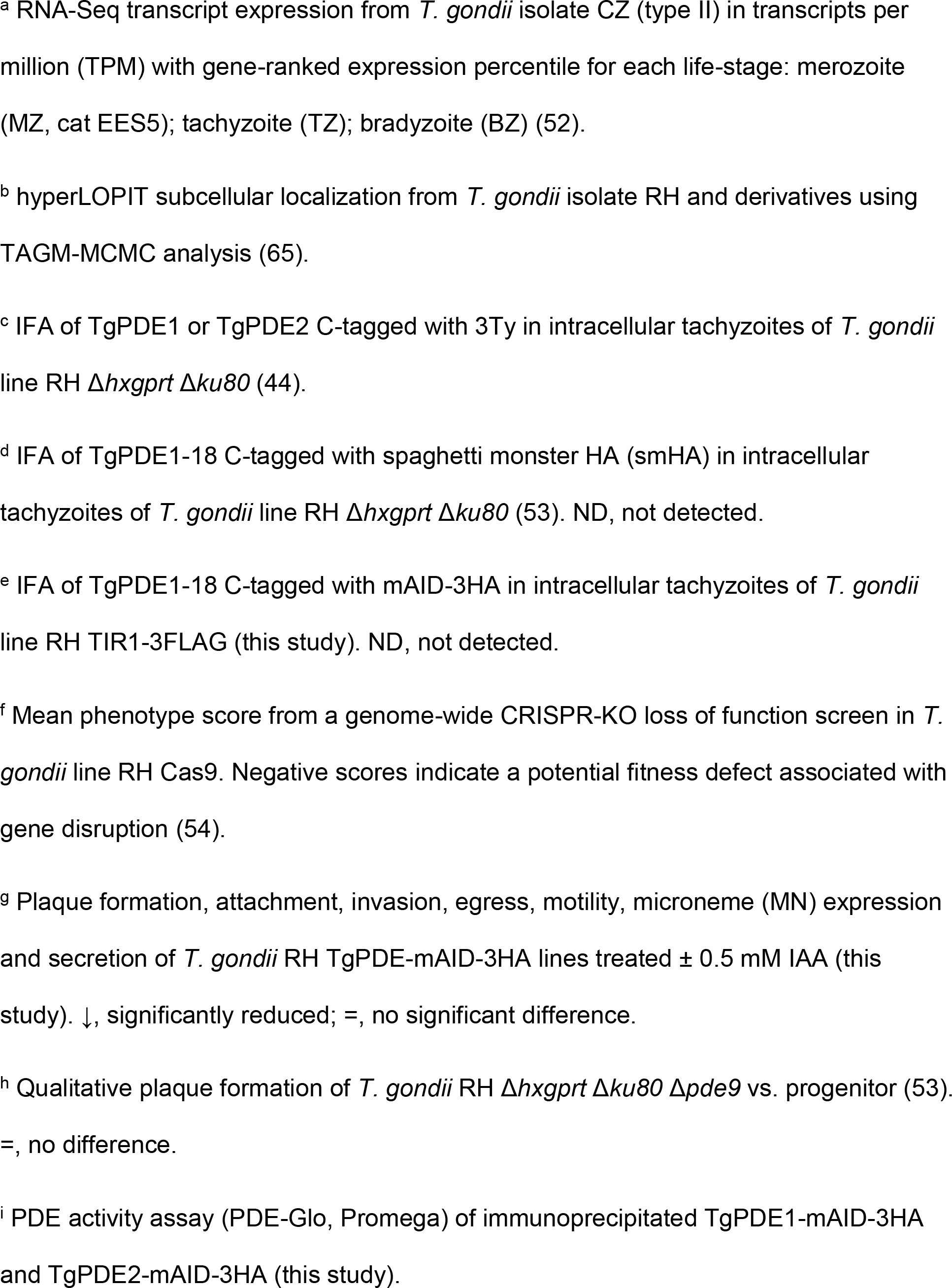
Summary of Toxoplasma PDE expression, localization, and function.

| Gene ID | Gene product | Mass (kDa) | Transcript expression TPM (%ile)<sup>a</sup> | Tachyzoite hyperLOPIT (probability)<sup>b</sup> | Tachyzoite distribution by IFA | Tachyzoite fitness score<sup>f</sup> | Tachyzoite cKD/ KO phenotypes | cNMP substrate |
| --- | --- | --- | --- | --- | --- | --- | --- | --- |
| TGGT1_202540 | TgPDE1 | 182.5 | MZ 27.85 (56.3)<br>TZ 70.84 (73.52)<br>BZ 34.79 (64.93) | Golgi (0.75)<br>PM – integral (0.25) | PM<sup>cde</sup><br>Cytomembranes<sup>cde</sup> | -2.43 | ↓Plaquing<sup>g</sup><br>=Replication<sup>g</sup><br>=Attachment<sup>g</sup><br>=Invasion<sup>g</sup><br>=Egress<sup>g</sup><br>↓Motility<sup>g</sup><br>↓MN expression<sup>g</sup> | cGMP<sup>i</sup><br>cAMP<sup>i</sup> |
| TGGT1_293000 | TgPDE2 | 243.9 | MZ 59.28 (75.98)<br>TZ 27.69 (49.57)<br>BZ 11.43 (41.04) | Nucleus – Chromatin (0.98) | ER<sup>cde</sup><br>Cytomembranes<sup>cde</sup> | -2.53 | ↓Plaquing<sup>g</sup><br>↓Replication<sup>g</sup><br>↓Attachment<sup>g</sup><br>↓Invasion<sup>g</sup><br>↓Egress<sup>g</sup><br>↓Motility<sup>g</sup><br>↓MN secretion<sup>g</sup> | cAMP<sup>i</sup> |
| TGGT1_233065 | TgPDE3 | 92.3 | MZ 30.45 (59.1)<br>TZ 0.34 (16.86)<br>BZ 0.83 (17.8) | - | ND<sup>de</sup> | -0.65 | =Plaquing<sup>g</sup> | - |
| TGGT1_229405 | TgPDE4 | 118.0 | MZ 73.14 (80.03)<br>TZ 2.21 (25.03)<br>BZ 23.25 (55.58) | - | ND<sup>de</sup> | -0.33 | =Plaquing<sup>g</sup> | - |
| TGGT1_220420 | TgPDE5 | 116.1 | MZ 2.06 (18.11)<br>TZ 4.56 (27.93)<br>BZ 12.74 (43) | - | ER<sup>d</sup><br>Cytomembranes<sup>d</sup><br>Mitochondrion<sup>e</sup> | 1.23 | ↓Plaquing<sup>g</sup> | - |
| TGGT1_266920 | TgPDE6 | 119.9 | MZ 3.42 (19.96)<br>TZ 399.87 (95.27)<br>BZ 538.83 (96.7) | PM - peripheral 2 (1) | Cytomembranes<sup>de</sup><br>Nucleus<sup>e</sup> | 1.63 | =Plaquing<sup>g</sup> | - |
| TGGT1_280410 | TgPDE7 | 121.6 | MZ 7.21 (25.9)<br>TZ 36.41 (56.66)<br>BZ 20.05 (52.25) | Golgi (0.75)<br>PM – integral (0.25) | PM<sup>de</sup><br>Cytomembranes<sup>d</sup> | 0.6 | =Plaquing<sup>g</sup> | - |
| TGGT1_318675 | TgPDE8 | 126.6 | MZ 32.06 (60.73)<br>TZ 3.36 (26.5)<br>BZ 13.48 (44.11) | - | Apical PM<sup>d</sup><br>Cytomembranes<sup>d</sup><br>ND<sup>e</sup> | 0.19 | =Plaquing<sup>g</sup> | cGMP<sup>i</sup><br>cAMP<sup>i</sup> |
| TGGT1_241880 | TgPDE9 | 142.5 | MZ 13.83 (37.77)<br>TZ 84.01 (77.12)<br>BZ 52.08 (72.88) | Apical 1 (0.51)<br>PM – peripheral 2 (0.34)<br>Golgi (0.14) | Apical PM<sup>de</sup> | 0.52 | ↓Plaquing<sup>g</sup><br>=Plaquing<sup>h</sup> | cGMP<sup>i</sup><br>cAMP<sup>i</sup> |
| TGGT1_272650 | TgPDE10 | 141.3 | MZ 5.56 (23.22)<br>TZ 53.07 (66.75)<br>BZ 15.13 (46.39) | PM – integral (0.5)<br>Golgi (0.5) | PM<sup>de</sup><br>Cytomembranes<sup>de</sup> | -0.37 | =Plaquing<sup>g</sup> | - |
| TGGT1_224840 | TgPDE11 | 146.8 | MZ 8.85 (28.99)<br>TZ 30.83 (52.57)<br>BZ 21.76 (53.9) | - | ER<sup>e</sup><br>Cytomembranes<sup>e</sup><br>ND<sup>d</sup> | 0.4 | =Plaquing<sup>g</sup> | - |
| TGGT1_310520 | TgPDE12 | 174.0 | MZ 13.59 (37.29)<br>TZ 7.74 (30.61)<br>BZ 24.58 (56.84) | - | Cytomembranes<sup>d</sup><br>ND<sup>e</sup> | -0.06 | =Plaquing<sup>g</sup> | - |
| TGGT1_257080 | TgPDE13 | 180.3 | MZ 16.77 (42.59)<br>TZ 55.8 (67.98)<br>BZ 22.97 (55.26) | ER (0.56)<br>Golgi (0.42) | ER<sup>e</sup><br>Basal<sup>d</sup><br>Cytomembranes<sup>de</sup> | 0.55 | =Plaquing<sup>g</sup> | - |
| TGGT1_228500 | TgPDE14 | 35.2 | MZ 42.17 (68.08)<br>TZ 0.65 (19.65)<br>BZ 2.64 (23.77) | - | ND<sup>de</sup> | -0.33 | =Plaquing<sup>g</sup> | - |
| TGGT1_233040 | TgPDE15 | 194.9 | MZ 39.65 (66.4)<br>TZ 0.79 (20.66)<br>BZ 7.3 (33.65) | - | ND<sup>de</sup> | 1.22 | =Plaquing<sup>g</sup> | - |
| TGGT1_258508 | TgPDE16 | 68.4 | MZ 146.2 (89.36)<br>TZ 2.37 (25.34)<br>BZ 56.46 (74.36) | - | ND<sup>de</sup> | -1.19 | =Plaquing<sup>g</sup> | - |
| TGGT1_257945 | TgPDE17 | 254.1 | MZ 226.54 (92.59)<br>TZ 25.37 (47.43)<br>BZ 24.5 (56.79) | - | ND<sup>de</sup> | -1.81 | =Plaquing<sup>g</sup> | - |
| TGGT1_226755 | TgPDE18 | 381.2 | MZ 27.59 (56.03)<br>TZ 6.01 (29.32)<br>BZ 0.46 (15.73) | - | ER<sup>de</sup><br>Cytomembranes<sup>de</sup> | -1.15 | =Plaquing<sup>g</sup> | - |

Studies of PDE function in many apicomplexans have been hindered by the sheer number of PDEs encoded in their genomes. This is especially true for coccidian apicomplexans like *Toxoplasma* that encode 18 PDE genes (Fig. 1). In contrast, other apicomplexan groups like hematozoa (i.e. malarial parasites and piroplasms) encode only 2-4 PDE genes per species. This discrepancy begs the question: why do coccidian apicomplexans need so many PDEs? We speculate that PDE expansion in coccidia meets the demands of their asexual (acute and chronic) and sexual life-stages, compartmentalized signaling, and serves as a redundancy safeguard for added control. Transcriptome-wide expression studies have indicated a life-stage dependence for the majority of TgPDEs (52). In support, 11 of 18 TgPDEs were recently shown to be expressed in tachyzoites by IFA and immunoblotting (53), which we were able to largely confirm independently here with disparate epitope tags (Fig. 3). Additional studies are needed to determine which TgPDEs control cyclic nucleotide turnover in other life-stages such as bradyzoites, merozoites, gametes, and sporozoites. We and others have also observed distinct subcellular distributions for TgPDEs (44, 53), indicating that they may act on separate pools of cyclic nucleotides for finely-tuned signaling.

Previously we showed that cGMP biosynthesis by TgGC and subsequent TgPKG activation occurs at the plasma membrane (21, 33). Therefore, cGMP-TgPDEs positioned at the plasma membrane would be well-positioned to turn off TgPKG signaling following invasion. The localization of cAMP signaling is more complex and less centralized in *Toxoplasma* (4 TgACs, 2 TgPKArs, 3 TgPKAcs) (21, 44, 45, 50, 63), which may require compartmentalized TgPDEs to inactivate cAMP signaling for egress or development. Overall, it appears beneficial for *Toxoplasma* to have life-stage-dependent PDEs distributed throughout the cell to prevent or inactivate unwanted cyclic nucleotide signaling.

Despite expressing at least 11 TgPDEs in tachyzoites, we were able to identify several TgPDEs that individually contribute to tachyzoite fitness (Fig. 4). It should be noted that we could not confirm knockdown of TgPDEs expressed below the limit of protein detection with our system, which could lead to an underestimation of fitness-conferring TgPDEs. Conditional depletion of TgPDE5 and TgPDE9 modestly reduced the size of plaques formed on HFF monolayers, indicating that they partially contribute to tachyzoite fitness. TgPDE9 was previously determined to be capable of degrading cAMP and cGMP, but was deemed dispensable by knockout in tachyzoites (53). Given the time it takes to generate and maintain knockouts, compensatory suppressor mutations could emerge and mask knockout phenotypes. Since TgPDE8 has the same localization and substrate preference as TgPDE9 (53), upregulation of TgPDE8 could feasibly suppress phenotypes associated with Δ*Tgpde9*. The speed of TgPDE9-mAID-3HA knockdown is unlikely to allow for suppressors, revealing TgPDE9’s true contribution to tachyzoite fitness. With loss of TgPDE1, we observed a slight reduction in plaque numbers and the plaques that did form were substantially smaller compared to vehicle treatment. The reduction in plaque numbers indicates that TgPDE1 is essential in a subset of parasites within a population of knockdown mutants. TgPDE2 depletion elicited the most severe plaquing defects, reducing plaque size by ∼95%.

These data indicate that TgPDE1 and TgPDE2 are indispensable and non-redundant for tachyzoite fitness, which supports the hypothesis that TgPDE1 and TgPDE2 are refractory to deletion (44, 54).

To understand how TgPDE1 and TgPDE2 contribute to tachyzoite fitness, we first tested whether either had phosphodiesterase activity. Attempts to generate highly-active recombinant TgPDE catalytic domain fragments were unsuccessful (Fig. S2-4). We suspect that the recombinant proteins were not adequately refolded following solubilization of inclusion body preparations and will require further optimization. It is also possible that some TgPDEs are simply catalytically inactive or hydrolyze unknown substrates. As an alternative approach, we performed activity assays on tagged TgPDEs immunoprecipitated from *Toxoplasma* lysates (Fig. 5). TgPDE1 was capable of degrading cGMP and cAMP, while TgPDE2 was cAMP-specific. It is reasonable for TgPDE1 and TgPDE2 to have different substrate preferences since they are both required for tachyzoite fitness and likely serve non-redundant roles. Our data suggests that TgPDE2 is primarily responsible for cAMP degradation in tachyzoites. Therefore, we speculate that TgPDE1 is primarily responsible for cGMP turnover in tachyzoites, with minor contributions from TgPDE8 and TgPDE9 that are also dual-specific PDEs (53), and possibly TgPDE5. It is not known whether dual-specific PDEs act selectively in *Toxoplasma*, but may be important given the mutually exclusive nature of cGMP and cAMP signaling in *Toxoplasma* (12, 15). A combinatorial knockout/knockdown approach is needed to directly address functional redundancy within the TgPDE family.

To ensure that the phosphodiesterase activities we observed for TgPDE1 and TgPDE2 were not artifacts of unexpected interactions with other TgPDEs, we analyzed immunoprecipitated TgPDE1 and TgPDE2 fractions with and without chemical cross- linking by LC-MS/MS (Fig. S5, Table S4, S5). No other host or *Toxoplasma* PDEs co-immunoprecipitated with TgPDE1 or TgPDE2, indicating that TgPDE1 and TgPDE2 were solely responsible for the cyclic nucleotide degradation observed *ex vivo*.

However, we did detect unique interactors for TgPDE1 and TgPDE2 that may regulate their activities through proteolysis, localization, or phosphorylation. Identifying domains and modifications that control TgPDE1 and TgPDE2 activity is critical to understand the mechanisms by which they regulate cyclic nucleotide turnover for lytic growth.

To further dissect and distinguish TgPDE1 and TgPDE2 function, we tested whether they control specific steps or processes within the tachyzoite lytic cycle. We determined that TgPDE1 is important for maintaining proper microneme levels (Fig. 6G) required for extracellular motility (Fig. 6F). Depletion of TgPDE1 may reduce microneme expression through premature microneme secretion or a general fitness defect. Since depletion of TgPDE1 reduced plaque area by 58% without directly diminishing invasion, replication, or egress (Fig. 4), we suspect that these parasites have difficulties spreading to new cells due to decreased motility and/or extracellular survival. Depletion of TgPDE2 was highly pleotropic, suppressing replication, invasion, egress, motility, and microneme secretion (Fig. 6). The cumulative perturbations of each step in the lytic cycle explains the fewer and 95% smaller plaques observed (Fig. 4). Since TgPDE2 functions as a cAMP-specific PDE (Fig. 5C), we speculate that its depletion elevates cAMP throughout the lytic cycle causing aberrant activation of TgPKAc1, which is known to antagonize motility and egress normally, but can also cause replication defects when overexpressed or hyperactivated (44, 45).

Altogether, our investigation of the 18-member TgPDE family revealed that TgPDE1 and TgPDE2 are the primary phosphodiesterases in *Toxoplasma* tachyzoites. We determined that TgPDE1 is a dual-specific PDE but we hypothesize that it primarily degrades cGMP in *Toxoplasma*. To our knowledge, this work also designates TgPDE2 as the only known cAMP-specific PDE in Apicomplexa. We propose a two-part working model in which 1) TgPDE1 is activated following *Toxoplasma* invasion to turn off cGMP-stimulated motility for replication and 2) TgPDE2 is activated following *Toxoplasma* replication to turn off cAMP-signaling, which inhibits motility, for egress and cell-to-cell transmission. This work enables future studies aimed at determining how TgPDE1 and TgPDE2 are regulated and cooperate to control cyclic nucleotide turnover and lytic growth. Furthermore, due to their conservation, TgPDE1 and TgPDE2 will serve as models for coccidian dual-specific and cAMP-specific PDEs, respectively.

## Materials and Methods

### Parasite and host cell culture

*T. gondii* tachyzoites were maintained in human foreskin fibroblasts (HFF) monolayers at 37°C, 5% CO2 in D10 medium: Dulbecco’s Modified Eagle’s Medium (Gibco) supplemented with 10% fetal bovine serum (Gibco), 10 mM glutamine (Gibco) and 10 μg/ml gentamicin (Gibco). Similarly, D3 medium containing 3% FBS was also used for parasite cultures where indicated. Cell lines were routinely assessed for *Mycoplasma* contamination by PCR using a Myco-Sniff Mycoplasma PCR Detection Kit (MP Biomedicals). All parasite lines used and generated in this study are listed in Table S1.

### Antibodies

Mouse anti-HA.11 (Clone 16B12) and Mouse anti-HIS tag (Clone J099B12) were purchased from BioLegend. Rat anti-HA (Clone 3F10) was purchased from Roche. Rabbit anti-TgSAG1 antibodies was provided by John Boothroyd, Ph.D. (Stanford University), rat anti-TgSAG1 was provided by Vernon Carruthers, Ph.D. (University of Michigan), and rabbit anti-TgGRA7 was provided by David Sibley, Ph.D (Washington University in St. Louis). Goat secondary antibodies conjugated to infrared (IR) dyes and Alexa Fluor dyes were purchased from LI-COR and Invitrogen, respectively.

### Plasmids

All plasmids generated in this study were created by Q5 Site-Directed Mutagenesis (New England Biolabs) of existing plasmids or HiFi Gibson Assembly (New England Biolabs) of linear dsDNA fragments. Plasmid sequences were confirmed by Sanger sequencing (Genewiz) and mapped using SnapGene v5.2.4 (GSL Biotech). All plasmids used in this study are listed in Table S2.

### Primers and PCR

All synthetic ssDNA and dsDNA oligonucleotides were synthesized by Integrated DNA Technologies and are listed in Table S3. Protoscript II Reverse Transcriptase (New England Biolabs) was used to make cDNA PCR templates from parasite RNA (*Toxoplasma* GT1; *Plasmodium falciparum* NF54) extracted with a Monarch Total RNA Miniprep Kit (New England Biolabs). Genomic DNA (*Toxoplasma* RH derivatives) was extracted for PCR using a Monarch Genomic DNA Purification Kit (New England Biolabs). Q5 Polymerase (New England Biolabs) was used for cloning and tagging amplicon PCRs. Taq polymerase (New England Biolabs) was used for diagnostic PCRs.

### Sequence analysis

Annotated genomic, transcript, and protein sequences for each TgPDE from Type I reference strain GT1 were downloaded from https://www.toxodb.org/. Transmembrane domains within TgPDE protein sequences were predicted using TOPCONS, a consensus of OCTOPUS, PHILIUS, PolyPhobius, SCAMPI, and SPOCTOPUS algorithms (57). Protein domains and features were predicted using the NCBI conserved domain search against the CDD v3.19 – 58235 PSSMs database with an Expect Value threshold of 0.01 (55). The predicted transmembrane domains and features were drawn to scale (primary aa length) using Adobe Illustrator (Adobe, Inc.).

### Generation of TgPDE-mAID-3HA conditional knockdown mutants

RH TIR1-3FLAG tachyzoites were used for tagging TgPDEs with mAID-3HA as described (59). To tag each gene of interest (GOI), a p*SAG1*:*Cas9-GFP*, *U6*:*sgGOI 3’ UTR* plasmid (2-5 µg) was co-transfected with a corresponding 40 bp homology arm-flanked *mAID-3HA*, *HXGPRT* amplicon (2-5 µg) into 1x10^6^ RH TIR1-3FLAG tachyzoites in P3 buffer using a 4D-nucleofector (Lonza) with pulse code FI-158. Transfected parasites were selected with 25 μg/ml mycophenolic acid (Alfa Aesar) and 50 μg/ml xanthine (Alfa Aesar) in D10 medium. Following drug selection, clones were isolated by limiting dilution and epitope tags were confirmed by diagnostic PCR of genomic DNA.

### Knockdown of TgPDE-mAID-3HA protein fusions

Knockdowns were performed as described (59). RH TIR1-3FLAG or RH TgPDE-mAID-3HA tachyzoites cultivated in HFFs in D10 medium were treated with 0.5 mM 3-indoleacetic acid (auxin; IAA) (Sigma Aldrich) prepared in 100% ethanol (Pharmco) or vehicle alone (0.0789% w/v ethanol final concentration) and incubated at 37°C, 5% CO2 prior to protein detection and/or phenotypic analysis.

### Detection of mAID-3HA tagged proteins by indirect immunofluorescence microscopy

Tachyzoite-infected HFF monolayers grown on 12 mm #1 glass coverslips (Electron Microscopy Sciences) were fixed with 4% formaldehyde (Polysciences) in phosphate buffered saline (PBS), permeabilized with 0.1% Triton X-100 (MP Biomedicals), blocked with 10% normal goat serum (Gibco) in PBS, labeled with mouse anti-HA (1:1000), and then washed three times with PBS. Antibody-labeled proteins were fluorescently labeled with goat anti-mouse IgG-AF488 (1:2000), washed five times with PBS, rinsed with water, and mounted on 25x75x1 mm Superfrost Plus glass slides (VWR) with Prolong Gold (Invitrogen). Wide-field images were captured and analyzed with a 100x oil objective on an Axioskop 2 MOT Plus wide-field fluorescence microscope (Carl Zeiss, Inc.) running AxioVision LE64 software (Carl Zeiss, Inc.).

### SDS-PAGE and immunoblotting

For routine detection of proteins, tachyzoite pellets were lysed in an equal volume of 2x Laemmli buffer (64) containing 20 mM dithiothreitol (DTT) (Thermo Fisher Scientific). All other protein samples were mixed 4:1 with 5x Laemmli buffer containing 50 mM DTT. Proteins were separated on 4-20% TGX polyacrylamide gels (Bio-Rad) or 4-20% TGX Stain-Free polyacrylamide gels (Bio-Rad) by SDS-PAGE. Total protein in polyacrylamide gels was detected using a ChemiDoc MP Imaging System (Bio-Rad) using Stain-Free (Bio-Rad) or Oriole (Bio-Rad) stains, as indicated in the figure legends. For immunoblotting, proteins separated by SDS-PAGE in polyacrylamide gels were wet-blotted onto nitrocellulose membranes. The membranes were rinsed with PBS containing 0.1% Tween-20 (PBS-T), and then blocked with PBS-T containing 5% (w/v) fat-free powered milk (blocking buffer). Membranes were probed with primary antibodies diluted in blocking buffer, then washed three times with PBS-T. The membranes were then incubated in the dark with goat IR- dye secondary antibodies (LI-COR) diluted in blocking buffer with mixing, then washed five times with PBS-T. Membranes were imaged on a ChemiDoc MP Imaging System.

Gels and membrane blots were analyzed using Image Lab software (Bio-Rad). **Plaque assays.** Freshly-egressed tachyzoites (RH TIR1-3FLAG and RH PDE-mAID- 3HA lines) were harvested, counted on a hemocytometer, and inoculated (200 parasites/ well) onto confluent HFF monolayers growing in 6-well plates containing D10 medium. To determine the effect of human phosphodiesterase inhibitors on *Toxoplasma* fitness, wells were treated with 0.5 mM 3-isobutyl-1-methylxanthine (IBMX) (MP Biomedicals) prepared in 100% DMSO (Sigma Aldrich), 0.5 mM zaprinast (Tocris) prepared in 100% DMSO, or vehicle 0.1% DMSO. To determine the effect of conditional knockdown of a TgPDE on *Toxoplasma* fitness, wells were treated with 0.5 mM IAA or 0.0789% w/v ethanol (vehicle). In both experiments, plates were left undisturbed for 8 days in a 37°C, 5% CO2 incubator. Plaque formation was assessed by counting zones of clearance on EtOH-fixed, crystal violet-stained HFF monolayers. Each stained plate was scanned with high-definition digital scanner (Epson) to obtain representative images and for plaque area analysis. For plaque area measurements, the 10 centermost plaques from the representative wells shown were analyzed using ImageJ to define the plaque area in square pixels (pix^2^).

### Replication assays

HFFs were grown to confluency in D10 in 96-well clear bottom plates (Greiner Bio-One), then inoculated with 2x10^5^ tachyzoites in D3 per well and incubated at 37°C, 5% CO2. After 2 h, non-invaded parasites were washed away and wells were treated with vehicle (EtOH) or 0.5 mM IAA in D3 at 37°C, 5% CO2 to degrade the mAID-3HA fusion proteins. At 24 h post-infection, the monolayers were fixed with 4% formaldehyde in PBS for 10 min, permeabilized with cold methanol for 5 min, and blocked with 10% goat serum in PBS. Parasites and vacuoles were labeled with rat anti- TgSAG1 (1:5000) and rabbit anti-TgGRA7 (1:1000), respectively. Next, parasites were counterstained with Alexa Fluor-dye conjugated goat secondary antibodies (1:2000) and Hoechst 33258 dye (1:5000). Sixteen fields were imaged per replicate at 40X using a Cytation 5 plate-reading microscope running Gen5 software (BioTek Instruments). To determine the number of parasites per vacuole, at least 100 vacuoles were counted per replicate and averaged from N = 3 independent trials.

### Invasion assays

Parasites were grown in HFFs in D3 containing vehicle (EtOH) or 0.5 mM IAA for 16 hours at 37°C, 5% CO2 to degrade the mAID-3HA fusion proteins. 4x10^5^ of the treated parasites were inoculated onto fresh HFF monolayers ± EtOH or 0.5 mM IAA in 96-well clear bottom plates and allowed to settle for 10 min, then incubated at 37°C, 5% CO2 for 20 min to facilitate invasion. The media was removed and the wells were gently washed once with PBS to remove non-attached parasites. Invasion was stopped by 4% formaldehyde in PBS fixation for 10 min, and the monolayer was blocked with 10% normal goat serum in PBS. Extracellular parasites were labeled with rat anti-TgSAG1 (1:5000), then the monolayer was permeabilized with cold methanol for 5 min. Intracellular and extracellular parasites were labeled with rabbit anti-TgSAG1 (1:20000). Following washing, parasites were then labeled with Alexa Fluor dye- conjugated secondary antibodies (1:2000) and Hoechst 33258 dye (1:5000) and washed again. Sixteen fields were imaged per replicate at 40X using a Cytation 5 plate- reading microscope running Gen5 software from N = 2-3 trials. Parasites were gated based on red + green (extracellular) vs green (intracellular) differential staining.

### Egress assays

HFFs were grown to confluency in D10 in 96-well clear bottom plates, then inoculated with 5x10^3^ tachyzoites in fresh D3 per well and grown in a 37°C, 5% CO2 incubator. After 18 h, parasites were treated with vehicle (EtOH) or 0.5 mM IAA in D3 to deplete mAID-3HA-tagged proteins. At 40 h, parasites were stimulated with vehicle (DMSO), 0.5 mM zaprinast, or 2 µM A23187 for 5 min in a 37°C, 5% CO2 incubator. Next, monolayers were pre-fixed by adding 10% formaldehyde directly to the wells (1:3 dilution) for 5 min. The media formaldehyde mixture was removed and fixed with 4% formaldehyde in PBS for 10 min. The fixative was removed and cold 100% methanol was added to permeabilize the monolayers for 5 min, then washed with PBS and blocked with 10% normal goat serum in PBS. Parasites and vacuoles were labeled with rat anti-TgSAG1 (1:5000) and rabbit anti-TgGRA7 (1:1000), respectively. Next, parasites were counterstained with Alexa Fluor-dye conjugated goat secondary antibodies (1:2000) and Hoechst 33258 dye (1:5000). Twelve fields were imaged per replicate at 20X using a Cytation 5 plate-reading microscope running Gen5 software from N = 3 independent trials. At least 80 vacuoles were counted per replicate. Vacuoles containing ≥ 4 parasites were considered intact to exclude reinvaded parasites.

### Motility assays

Parasites were grown in HFFs in D3 containing vehicle (EtOH) or 0.5 mM IAA for 14 hours at 37°C, 5% CO2 to degrade the mAID-3HA fusion proteins, then harvested for motility assays on bovine serum albumin (BSA)-coated 96-well clear bottom plates. To coat the wells, 1% (w/v) BSA in extracellular (EC) buffer (142 mM NaCl, 1 mM MgCl2, 1.8 mM CaCl2, 5.6 mM D-glucose, 25 mM HEPES, pH 7.4) was added for 2 h at room temperature prior to the start of the assay. Freshly-purified treated parasites were counted and resuspended in EC buffer ± EtOH or 0.5 mM IAA. One-hundred µl of parasite suspension (10^5^ parasites) was added to empty BSA-coated wells and allowed to settle for 10 min. Following a 20 min incubation at 37°C, 5% CO2, parasites were fixed to the wells and immunolabeled with rat anti-TgSAG1 (1:5000) and goat anti-rat Alexa Fluor 488 (1:2000) to detect the parasites and their motility trails by IF microscopy. Twenty-five fields were imaged per replicate at 40X using a Cytation 5 plate-reading microscope running Gen5 software. At least 100 parasites per replicate were examined for TgSAG1-based motility trails and categorized by no trail, a short trail (< 1 body length), or a long trail (> 1 body length). Percentages of motile parasites were averaged from two replicates per treatment from 1 of N = 2 independent trials.

### Microneme secretion assays

Parasites were grown in HFFs in D3 containing vehicle (EtOH) or 0.5 mM IAA for 14 hours at 37°C, 5% CO2 to degrade the mAID-3HA fusion proteins, then harvested for microneme secretion assays. Treated parasites were counted and resuspended in EC buffer ± EtOH or 0.5 mM IAA at 10^8^/ml. A 100 µl aliquot of parasites (10^7^ parasites) was lysed with 20 µl 5x Laemmli buffer containing 50 mM DTT to determine the total protein content for subsequent analysis of secreted proteins by immunoblotting. Separately, 100 µl of parasites (10^7^ parasites) were incubated with vehicle (DMSO) or 0.5 mM zaprinast for 10 min at 37°C, 5% CO2 to facilitate microneme secretion, then chilled on ice. Parasites were centrifuged 2x at 800 x *g*, 4°C, 10 min to isolate parasite-free secreted fractions. Fifty µl of secreted fractions were mixed with 10 µl 5x Laemmli buffer containing 50 mM DTT. Twelve µl of total protein (diluted 1:10) and 12 µl of secreted fractions were resolved by SDS-PAGE (4- 20% TGX), transferred to nitrocellulose membranes, and blocked with 5% (w/v) milk PBS-T. Membranes were incubated with mouse anti-MIC2 (1:1000) and rabbit anti- TgGRA7 (1:1000), washed 3x with PBS-T, then incubated with IR-dye conjugated goat secondary antibodies (1:5000). Following 5x washes in PBS-T, membranes were imaged on a ChemiDoc MP Imaging System and analyzed using Image Lab software. Representative images were derived from 1 of N = 2 independent trials with similar outcomes.

### Immunoprecipitation of TgPDE1 and TgPDE2 for activity assays

Freshly-egressed tachyzoites (RH TIR1-3FLAG, RH PDE1-mAID-3HA, and RH PDE2-mAID-3HA) from HFF T150 cultures were scraped, syringe-lysed three times with a 25-gauge needle to liberate remaining intracellular parasites, and pelleted (800 x *g*, 4°C, 10 min). Pellets were washed with 30 ml cold PBS and counted on a hemocytometer, then pelleted again. Pellets were washed once more with 1 ml cold PBS, transferred to 2 ml tubes, then pelleted again. Pellets containing ∼ 1x10^8^ parasites were resuspended in 1 ml native lysis buffer (NLB) containing 10 mM K2HPO4, 150 mM NaCl, 5 mM EDTA, 5 mM EGTA, pH 7.4, 0.2% sodium deoxycholate, 1% Triton X-100, and 1X protease inhibitor cocktail with EDTA (Pierce), then incubated on ice for 30 minutes with mixing as described (53). Lysates were centrifuged at 14500 x *g*, 4°C, 20 min to remove insoluble material. The soluble supernatant was added to NLB-washed anti-HA magnetic beads (Pierce, 25 µl slurry per IP), and mixed at 4°C for 2 h. A Dynamag (Invitrogen) stand was used to separate beads from supernatant following IP, washing, and elution steps. The beads were washed once with 1 ml NLB, twice with 1 ml PDE storage buffer (40 mM Tris-HCl, 150 mM NaCl, pH 7.5) containing 0.05% Tween-20, and once with 0.2 ml PDE storage buffer. Immunoprecipitated proteins were eluted from beads with 0.2 ml PDE storage buffer containing 1 mg/ml HA-peptide (Thermo Fisher Scientific) with mixing at 37°C for 30 minutes. Bead-free elution fractions and other IP fractions were stored at -80°C until use.

### Immunoprecipitation of TgPDE1 and TgPDE2 for LC-MS/MS protein identification

Freshly-egressed tachyzoites (RH YFP-AID-3HA, RH PDE1-mAID-3HA, and RH PDE2- mAID-3HA) from HFF T150 cultures were scraped, syringe-lysed three times with a 25- ga needle to liberate remaining intracellular parasites, and pelleted (800 x *g,* 4°C, 10 min). Pellets were washed with 30 ml cold PBS, counted on a hemocytometer, then pelleted again. For crosslinked IP, parasite pellets were resuspended in cold 6 ml PBS containing 1% formaldehyde and incubated for 10 min. Following crosslinking, parasites were pelleted and resuspended in 6 ml PBS containing 125 mM glycine for 5 min to quench residual formaldehyde, then pelleted again. Fixed parasite pellets were then resuspended in 30 ml cold PBS. The non-crosslinked and crosslinked parasite suspensions were pelleted and resuspended in 6 ml NLB, syringe disrupted three times with a 25-ga needle, and then spun at 3200 x *g*, 4°C, 50 min to remove insoluble material. The 6 ml soluble lysate fractions were precleared with NLB-washed anti-c-Myc magnetic beads (Pierce, 50 µl slurry per IP) with mixing for 2 h at 4°C. The anti-c-Myc beads were removed by centrifugation (500 x *g*, 4°C, 10 min), and the precleared soluble lysates were incubated with pre-washed anti-HA magnetic beads (Pierce, 50 µl slurry per IP) overnight at 4°C with mixing. The anti-HA magnetic beads were centrifuged (500 x *g*, 4°C, 10 min) and transferred to 1.5 ml tubes for washing using a Dynamag stand. The beads were washed five times with 1 ml NLB and three times with 1 ml PBS. Dry beads and other IP fractions were stored at -80°C until use. Dry beads containing captured proteins were shipped on dry ice to the Proteomics & Metabolomics Facility at the Center for Biotechnology/University of Nebraska – Lincoln for liquid chromatography with tandem mass spectrometry (LC-MS/MS) protein identification. Specific LC-MS/MS analysis parameters are listed in Table S4 (native IP) and S5 (crosslinked IP), respectively.

### Expression and purification of 6HIS-SUMO-TgPDE^CAT^ proteins

SHuffle T7 Competent *E. coli* (New England Biolabs) were transformed with pET-*6HIS-SUMO- “Protein”* plasmids containing *TgPDE1-18*^CAT^, *PfPDEα*^CAT^, *PfPDEβ*^CAT^, or *HsGSDMD* fusions (listed in Table S2). Transformed *E. coli* starter cultures were grown overnight at 30°C in 50 ml Terrific Broth (TB) (Fisher Scientific) containing 50 µg/ml kanamycin with shaking. The cultures were split 1:5 in 250 ml TB containing kanamycin and grown at 30°C until reaching an OD600 between 0.6 and 0.8. The cultures were induced with 1 mM isopropylthio-β-galactoside (IPTG) (Life Technologies) and grown for 4 h at 30°C. The cells were centrifuged (12000 x *g*, 4°C, 20 min) and the cell pellets were stored at - 20°C overnight. The frozen pellets were resuspended in 25 ml cold homogenization buffer: PBS pH 7.4 containing 10 µl/ml Halt Protease Inhibitor Cocktail EDTA-free (Thermo Fisher Scientific) and 2 µl/ml benzonase (Sigma Aldrich). *E. coli* were homogenized using an Emusiflex-C5 (Avestin) with 6 cycles at 20,000 psi. The resulting homogenates were centrifuged (12000 x *g*, 4°C, 20 min) and the insoluble inclusion body fraction was collected (fraction determined to contain recombinant proteins) and solubilized in 4 ml PBS containing 10% (w/v) sarkosyl (MP Biomedicals) with rocking overnight at 4°C. The samples were diluted 1:10 in 40 ml PBS and insoluble material was removed by centrifugation (12000 x *g*, 4°C, 20 min). The soluble supernatants were added to 2 ml pre-equilibrated Ni-NTA slurry (Fisher Scientific), incubated for ∼5 h with mixing, then loaded onto to a 10 ml gravity column (Thermo Fisher Scientific) for washing and elution. The resins were washed twice with 40 ml recombinant wash buffer (40 mM Tris-HCl, pH 7.5, and 5 mM imidazole). The recombinant 6HIS-SUMO protein fusions were eluted with 3 ml recombinant elution buffer (40 mM Tris-HCl, pH 7.5, and 250 mM imidazole) and concentrated to 500 µl using Amicon Ultra 10 kDa Spin Columns (Millipore Sigma). The concentrated solutions underwent two buffer exchanges with 500 µl 40 mM Tris-HCl in 0.5 ml Amicon Ultra 10 kDa Spin Columns to reduce the imidazole concentration from 250 mM to ∼ 2.5 mM. The partially-purified recombinant proteins were analyzed for purity and identity by SDS-PAGE and immunoblotting with mouse anti-HIS tag, respectively. Protein concentrations were determined using the Rapid Gold BCA Protein Assay kit (Pierce) measured with a Cytation 5 plate imager running Gen5 software. The recombinant proteins were aliquoted and stored at -80°C until use.

### Phosphodiesterase activity assays

To assess the substrate specificity of TgPDEs, recombinant TgPDEs or immunoprecipitated TgPDEs were assayed using the PDE-Glo Phosphodiesterase Assay kit (Promega) according to the manufacturer’s instructions with a few modifications. In brief, cyclic nucleotides were diluted in Reaction Buffer (40 mM Tris-HCl pH 7.5, 50 mM MgCl2, 0.5 mg/ml BSA). On ice, 10 μl cAMP (2 or 0.2 µM) or cGMP (20 µM) were combined with 10 μl (1 μg) of purified recombinant 6HIS-SUMO- PDE^CAT^ protein or 10 μl immunoprecipitated eluate in triplicate to a 96-well non-skirted natural PCR plate (Midwest Scientific). Cyclic nucleotide standards were diluted further in Reaction Buffer, then 10 μl of standards were combined with 10 μl PDE storage buffer in triplicate in the same plate. To initiate the phosphodiesterase reaction, the plates were sealed and incubated at 37°C for 2 h in a T100 Thermal Cycler (Bio-Rad).

After incubation, 10 μl of the phosphodiesterase reaction was added to a 96-well half area flat bottom white polystyrene plate (Greiner) containing 5 μl Termination Buffer. Five μl of Detection Buffer was added to each reaction, covered with parafilm, then incubated at room temperature for 20 min. After incubation, 20 μl Kinase-Glo was added to the reactions, covered with parafilm and foil, and incubated at room temperature for 10 min. The resulting luminescence was quantified using a Cytation 5 plate imager running Gen5 software.

### Statistical analysis

Data was graphed and analyzed for statistical significance using GraphPad Prism v 9 (GraphPad Software) using a two-tailed Student’s T test for pairwise comparisons of normally distributed data. A two-way ANOVA with Tukey’s multiple comparisons test was used for comparing data from multiple treatment conditions with multiple outcomes or observations. Error bars represent the standard deviation or standard error of the mean as indicated in the figure legends. Differences between means were considered statistically significant when *P* was < 0.05 and designated with one or more asterisks above the graphed data. Non-significant differences between means (*P* > 0.05) were labeled with “ns” in the figures.

### Data availability

A merged Scaffold file (.sf3) of all LC-MS/MS data and annotated results is available upon request.

## Acknowledgements

We thank Sophie Alvarez, Ph.D. and Michael Naldrett, Ph.D. at the Proteomics & Metabolomics Facility (RRID:SCR_021314) at the Center for Biotechnology/University of Nebraska – Lincoln for LC-MS/MS analysis. The facility and instrumentation are supported by the Nebraska Research Initiative. We thank John Boothroyd, Ph.D. (Stanford University) and the *Toxoplasma* community for antibodies and other key reagents used in this study. We thank Josh Beck, Ph.D. (Iowa State University) for *Plasmodium falciparum* NF54 lysates. Special thanks to L. David Sibley, Ph.D. (Washington University in St. Louis) for professional guidance and support during the developmental stages of this project. This research was supported in part by National Institutes of Health NIAID grant # K22AI144035 to KMB.

---

**FIG S1.**
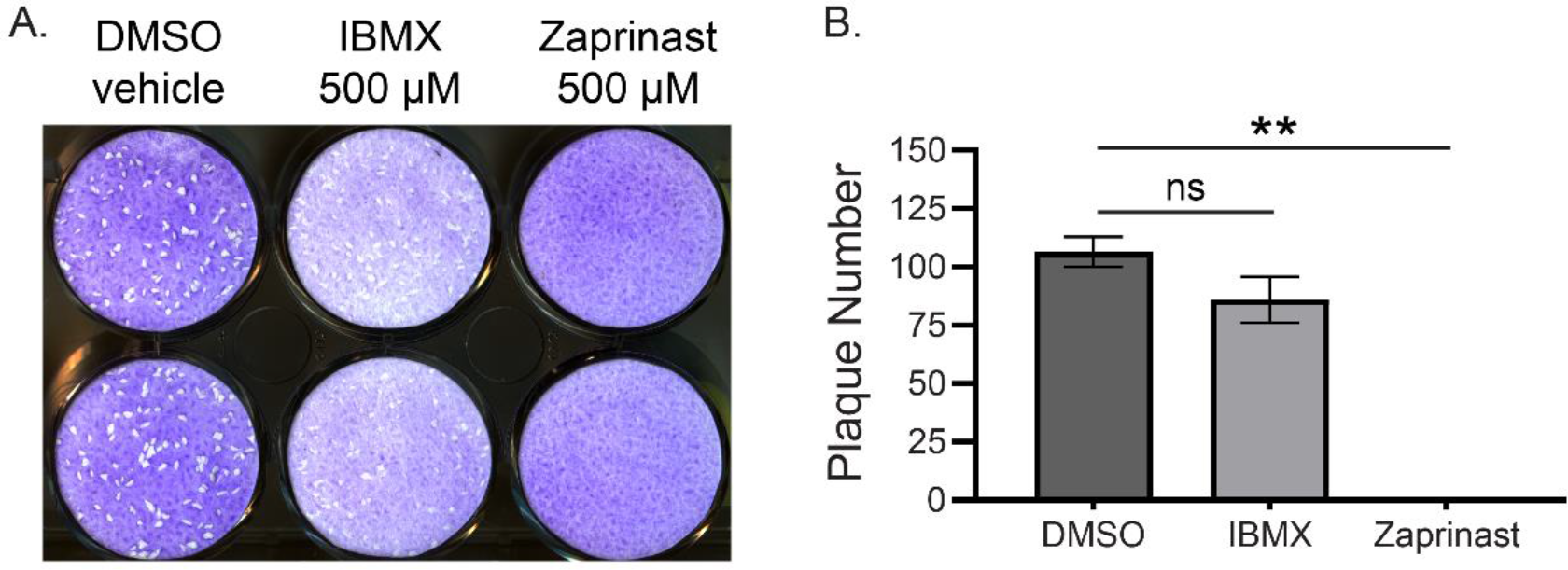
Pharmacological evidence for the importance of TgPDEs. (A) Plaques formed on HFF monolayers by 200 RH TIR1-3FLAG tachyzoites treated with vehicle (DMSO) or 0.5 mM PDE inhibitors (IBMX or zaprinast) for 8 days. (B) Quantification of plaques shown in A, mean ± SD (n = 2). Statistical significance was determined using an unpaired student’s t test comparing plaque formation in the presence of PDE inhibitors vs vehicle, \*\**P* ≤ 0.01.

**FIG S2.**
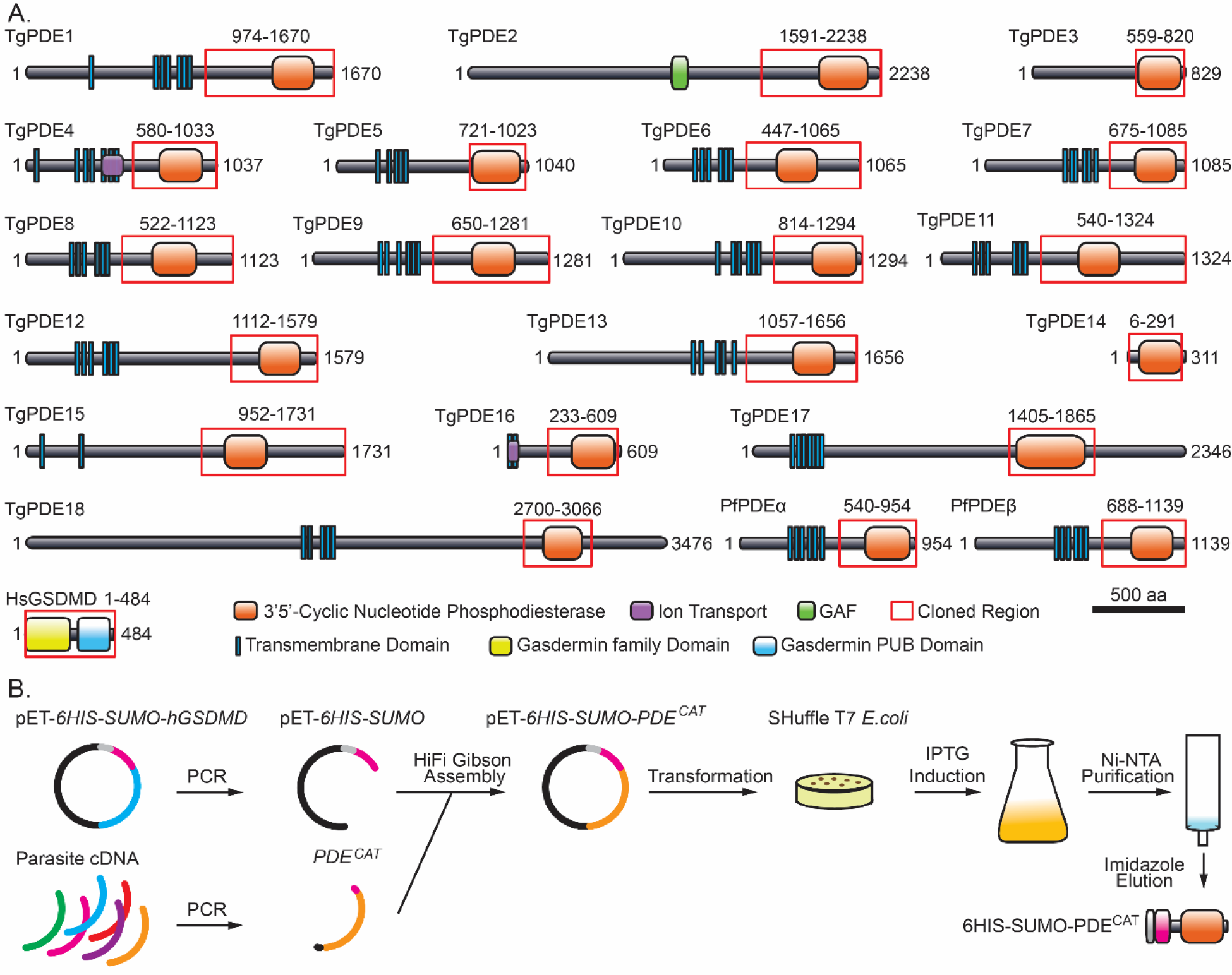
Strategy for production of recombinant TgPDE catalytic fragments. Related to FIG S3-4. (A) Cloned regions (outlined in red) for 18 TgPDEs, 2 PfPDEs (+ controls), and human gasdermin D (hGSDMD) protein (- control). (B) Cloning, transformation, induction, and purification of recombinant 6HIS-SUMO protein fusions in *E. coli*.

**FIG S3.**
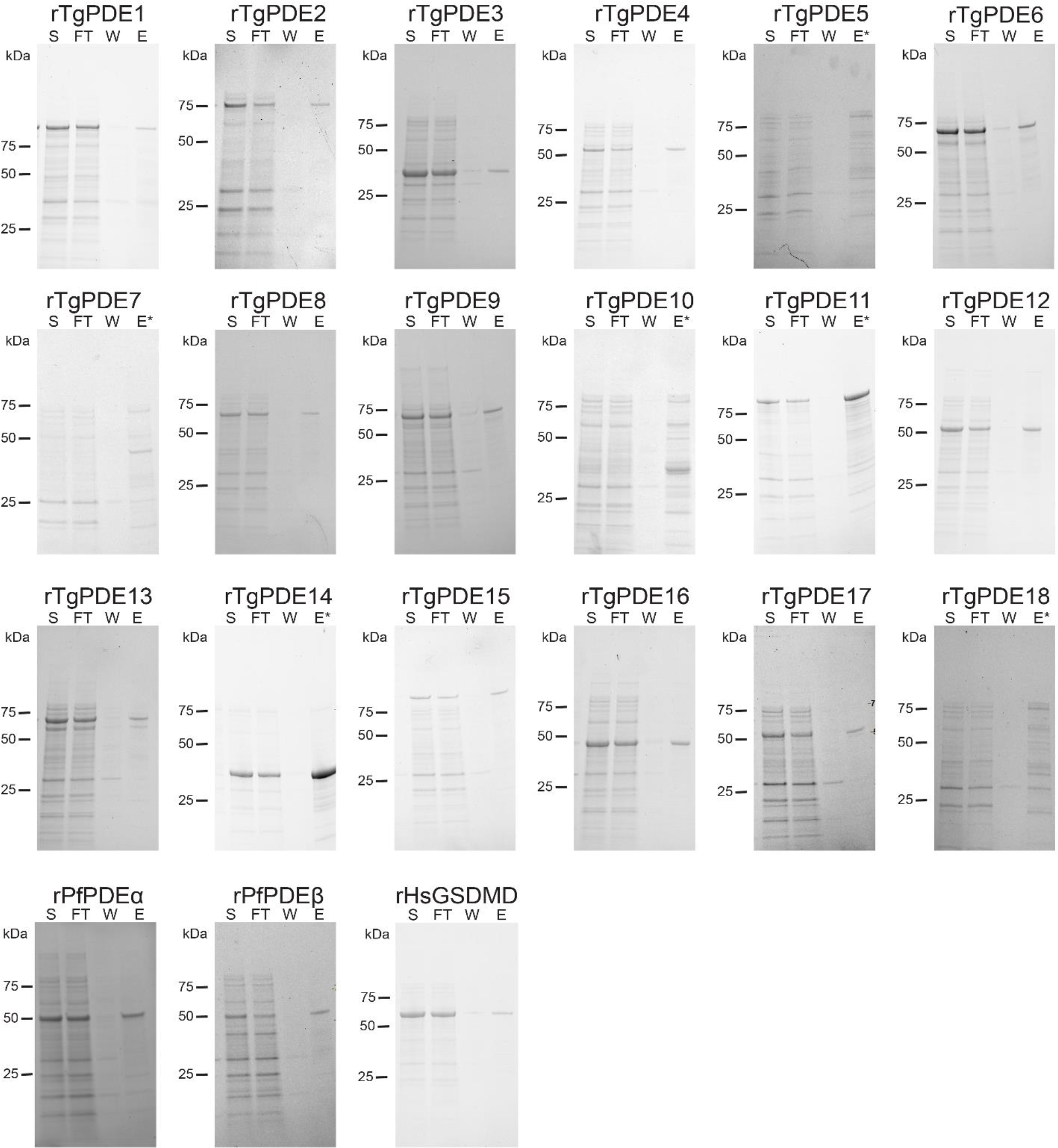
SDS-PAGE analysis of recombinant TgPDE catalytic fragments. Related to FIG S2, FIG S4. Homogenized inclusion body fractions from IPTG-induced *E. coli* expressing 6HIS-SUMO protein fusions were solubilized with sarkosyl and partially purified using immobilized metal affinity chromatography (Ni-NTA). Recombinant protein fractions from a representative experiment were resolved on a 4-20% SDS-PAGE gel containing total protein fluorescent dye for imaging. S, soluble lysate; FT, flowthrough; W, combined washes; E, 1:10 diluted imidazole eluate (recombinant protein); E*, undiluted imidazole eluate.

**FIG S4.**
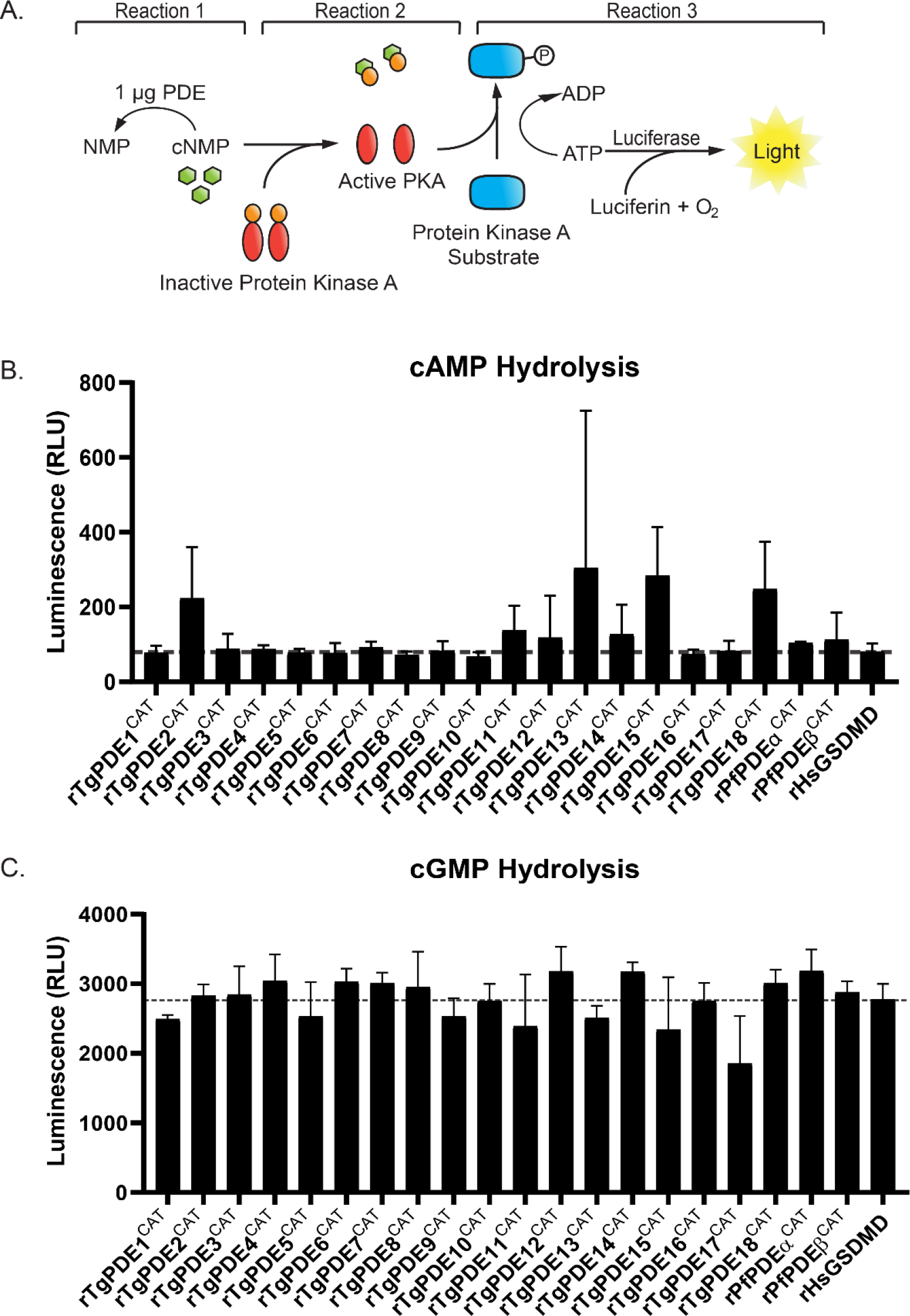
Phosphodiesterase activities of recombinant TgPDE catalytic fragments. Related to FIG S2-3. (A) Strategy to determine phosphodiesterase activity of recombinant PDEs (6HIS-SUMO fusions) using the PDE-Glo system where relative luminescence (RLU) is proportional to cyclic nucleotide hydrolysis. (B) Cyclic AMP phosphodiesterase activity of 1 µg recombinant proteins incubated 1:1 with 2 µM cAMP for 2 h at 37°C. (C) Cyclic GMP phosphodiesterase activity of 1 µg recombinant proteins incubated 1:1 with 20 µM cGMP for 2 h at 37°C. (B,C) Dotted lines represent the baseline activity threshold of the negative control, recombinant human gasdermin D (rHsGSDMD). A single trial of several similar trials is shown. Error bars indicate the standard deviation of 3 technical replicates.

**FIG S5.**
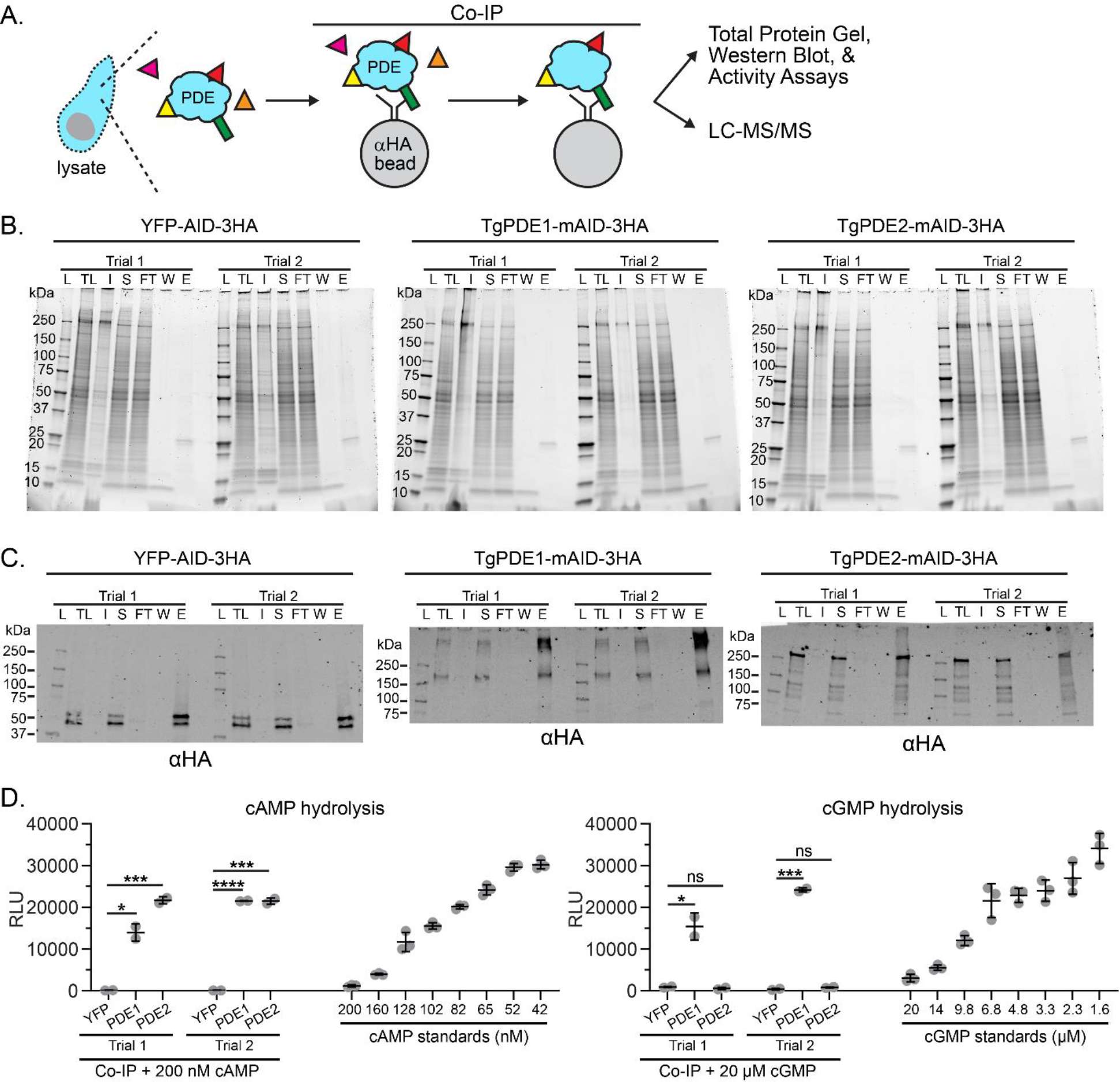
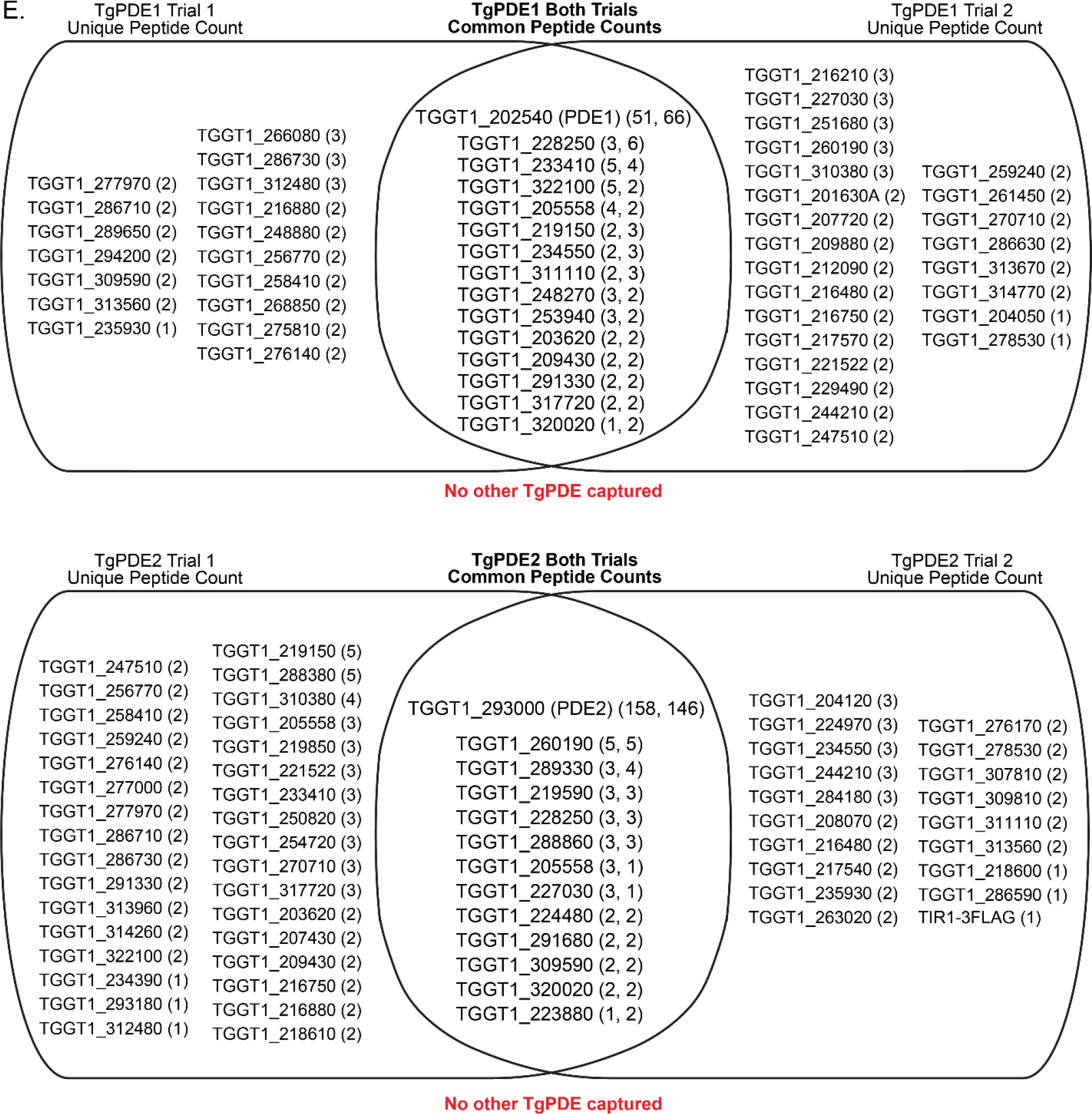
Protein interactomes of TgPDE1 and TgPDE2. (A) Schematic of co-immunoprecipitation of mAID-3HA fusions with anti-HA magnetic beads and downstream applications. (B) SDS-PAGE analysis of total protein in co-immunoprecipitation fractions using Oriole staining. L = ladder, TL = total lysate, I = insoluble lysate, S = soluble lysate, FT = flowthrough, W = combined washes, E = eluate. (C) Immunoblot analysis of co-immunoprecipitation fractions probed with rat anti- HA (1:1000) and goat anti-rat IRDye 800CW (1:5000). (D) Left, cAMP phosphodiesterase activity of immunoprecipitated elution fractions incubated 1:1 with 0.2 µM cAMP for 2h at 37°C. Standards shown were incubated 1:1 with PDE storage buffer for 2 h at 37°C. Right, cGMP phosphodiesterase activity of immunoprecipitated elution fractions incubated 1:1 with 20 µM cGMP for 2h at 37°C. Standards shown were incubated 1:1 with PDE storage buffer for 2 h at 37°C. Data represented as mean ± SD (n = 3) for each trial shown. Statistical significance was determined using an unpaired student’s t test comparing phosphodiesterase activity between PDE fractions and YFP (negative control). *P*: * ≤ 0.05, *** ≤ 0.001, **** ≤ 0.0001. (E) LC-MS/MS identification of protein interactors of TgPDE1 and TgPDE2 from two independent co- immunoprecipitation trials. Shown are the gene accessions with unique peptide counts for each protein identified (not found in YFP negative control). No other TgPDEs co- purified with TgPDE1 or TgPDE2.

**Table S1.**
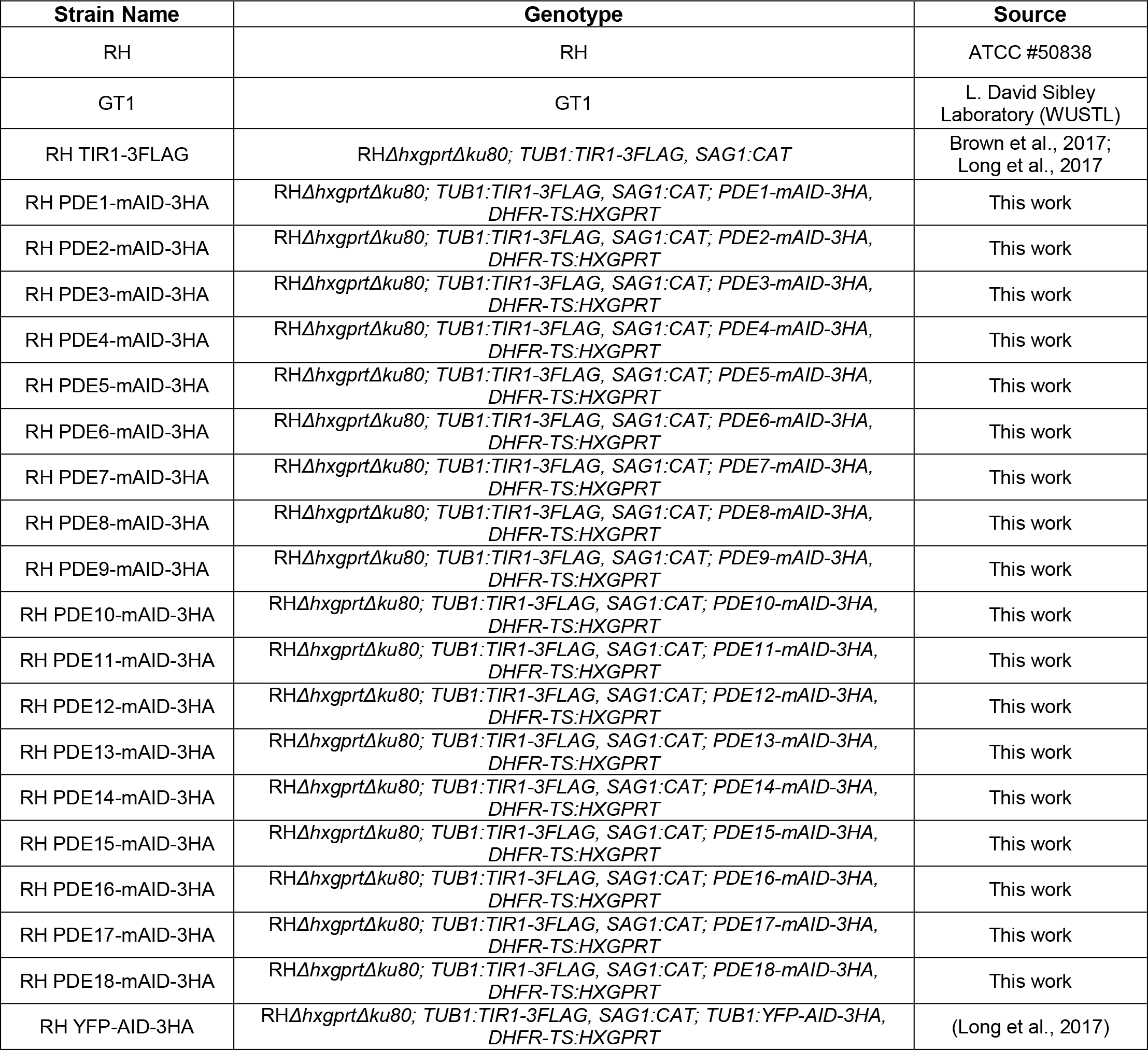
Toxoplasma strains used in this study.

**Table S2.**
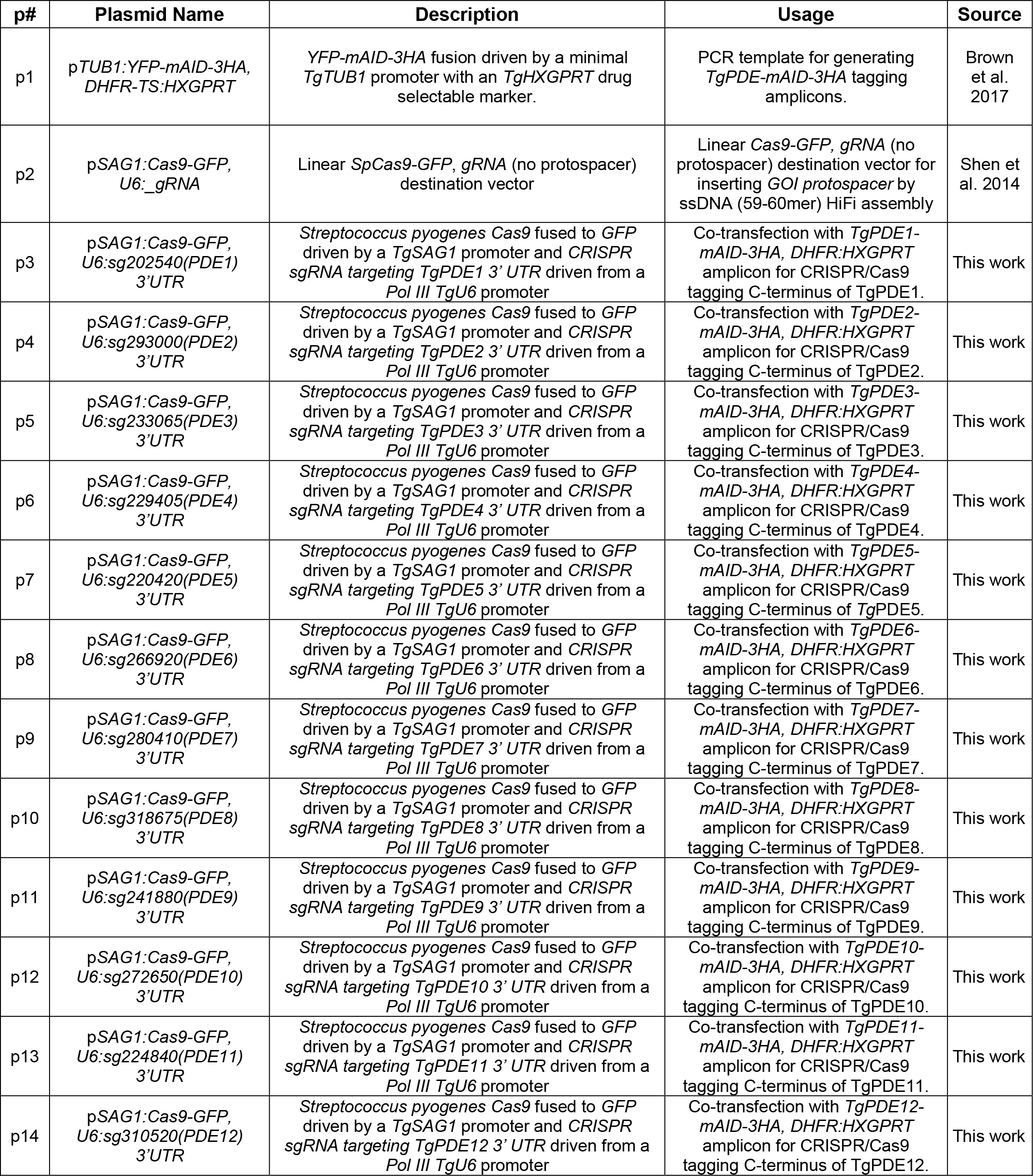
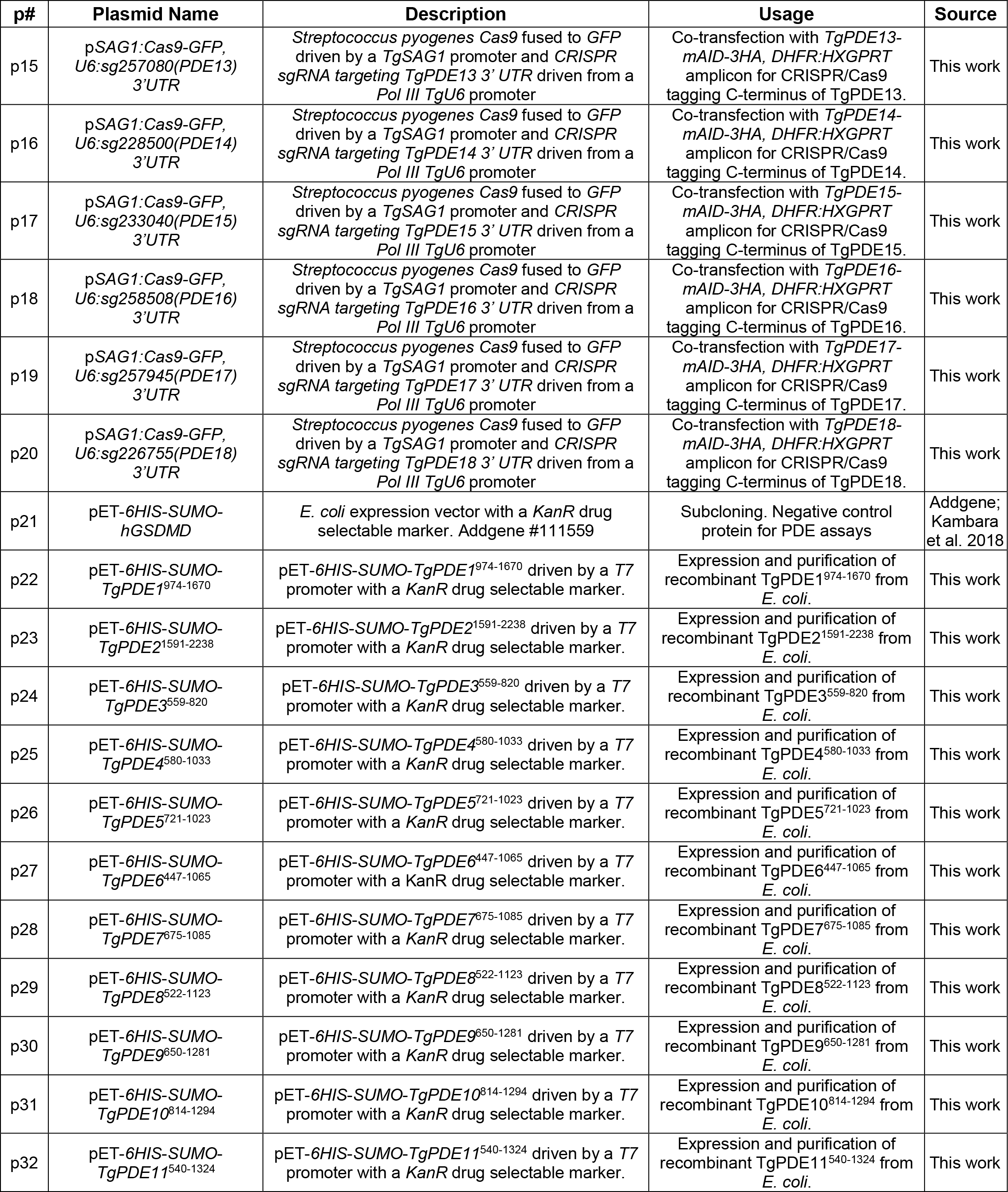
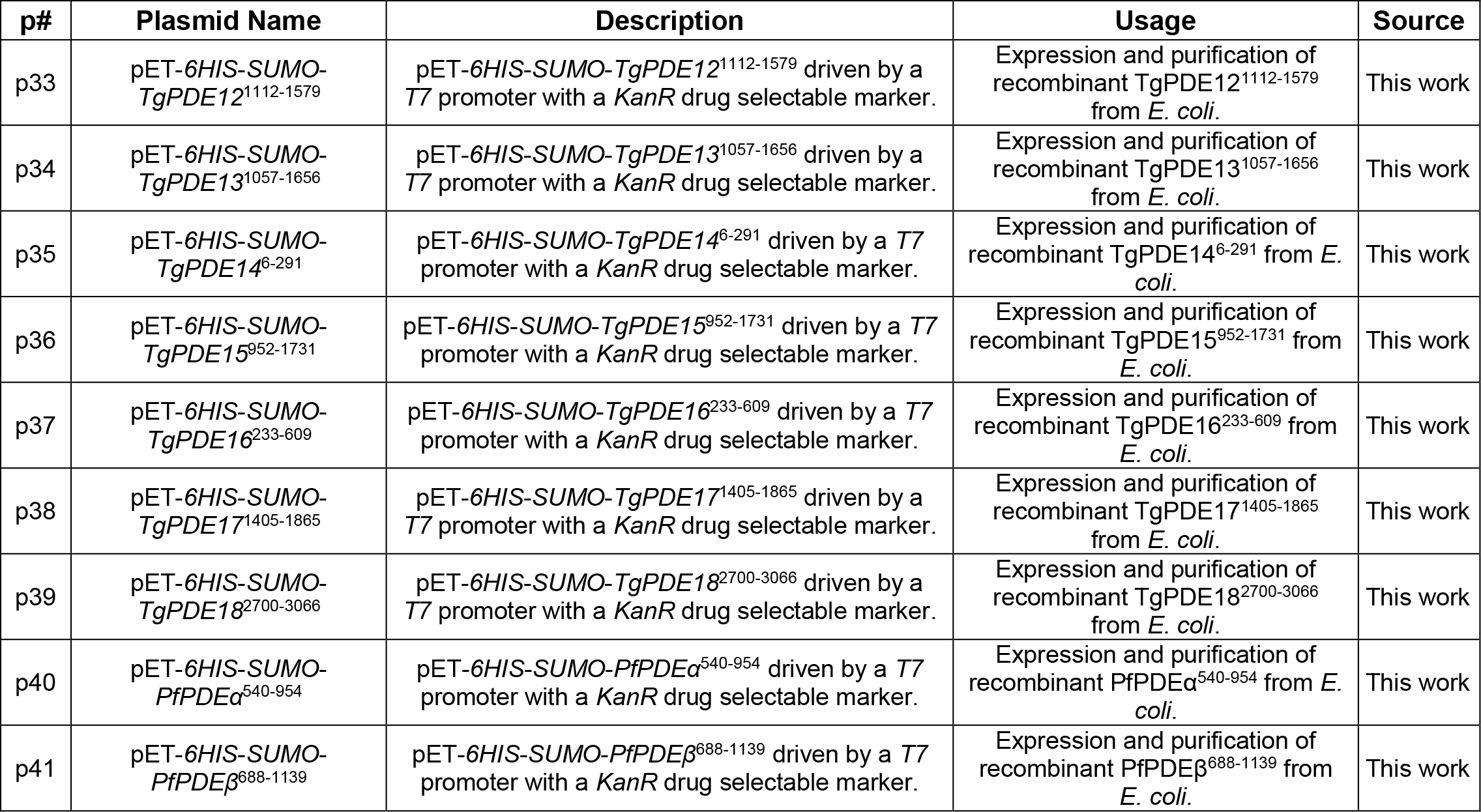
Plasmids used in this study.

**Table S3.**
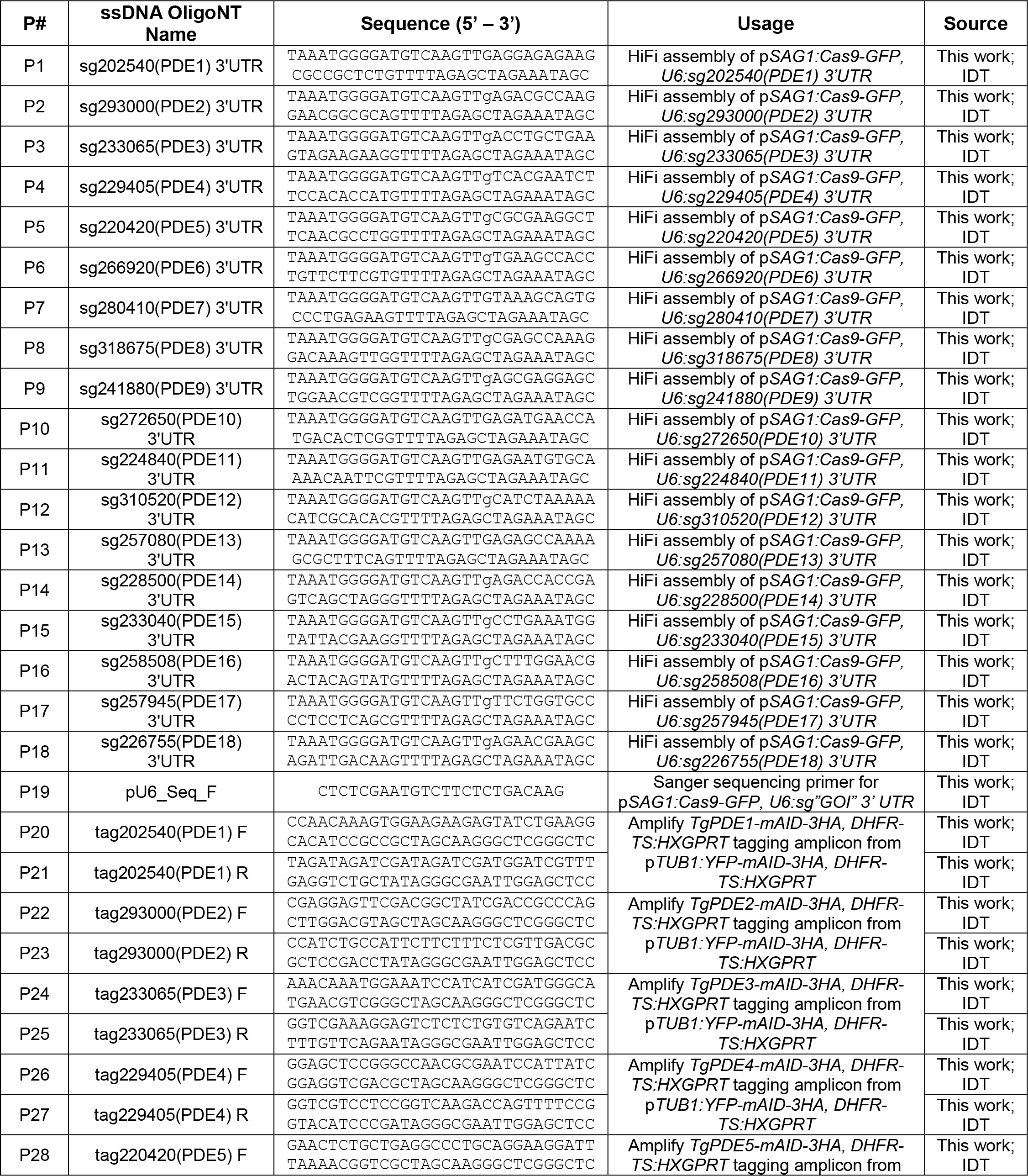
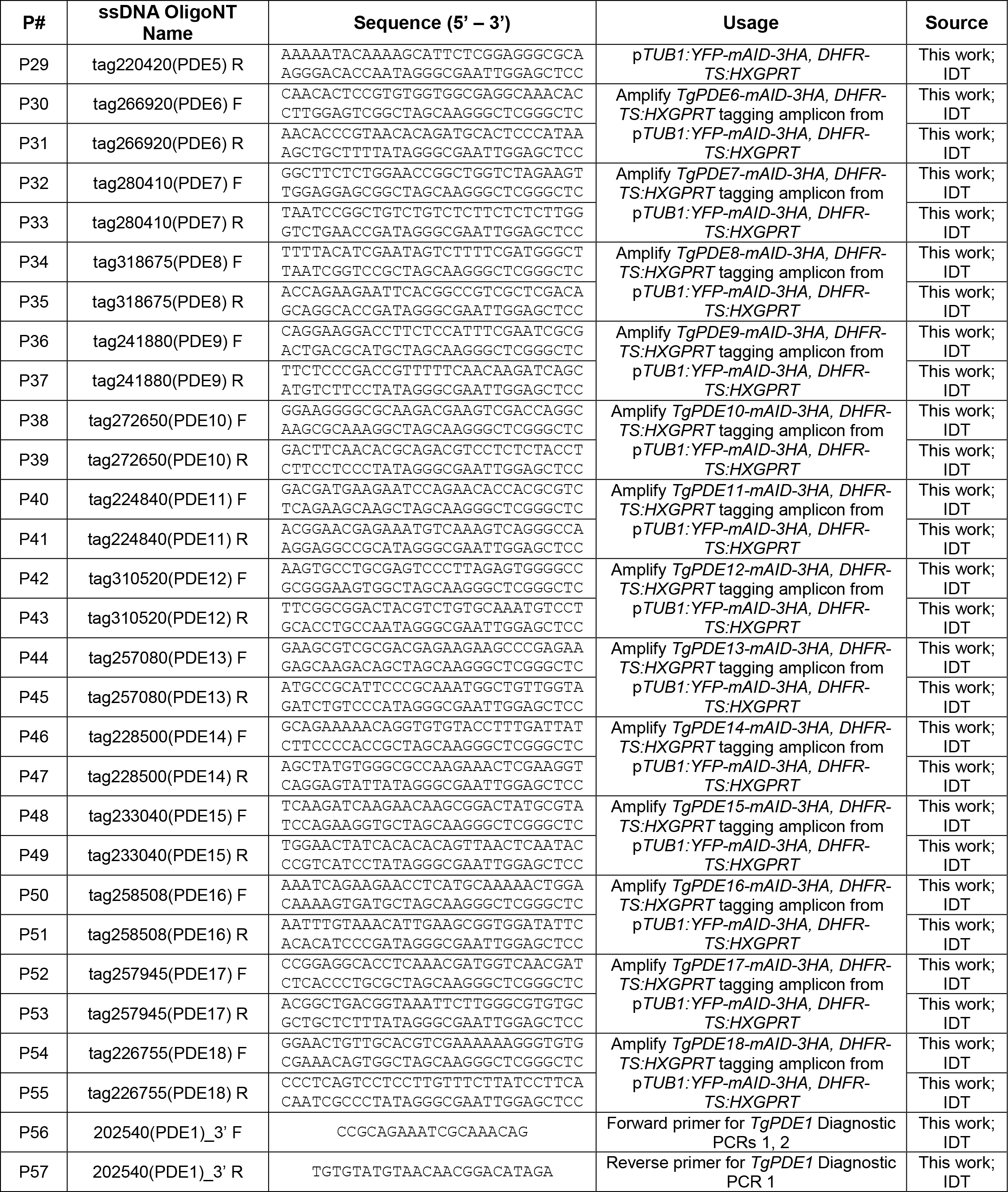
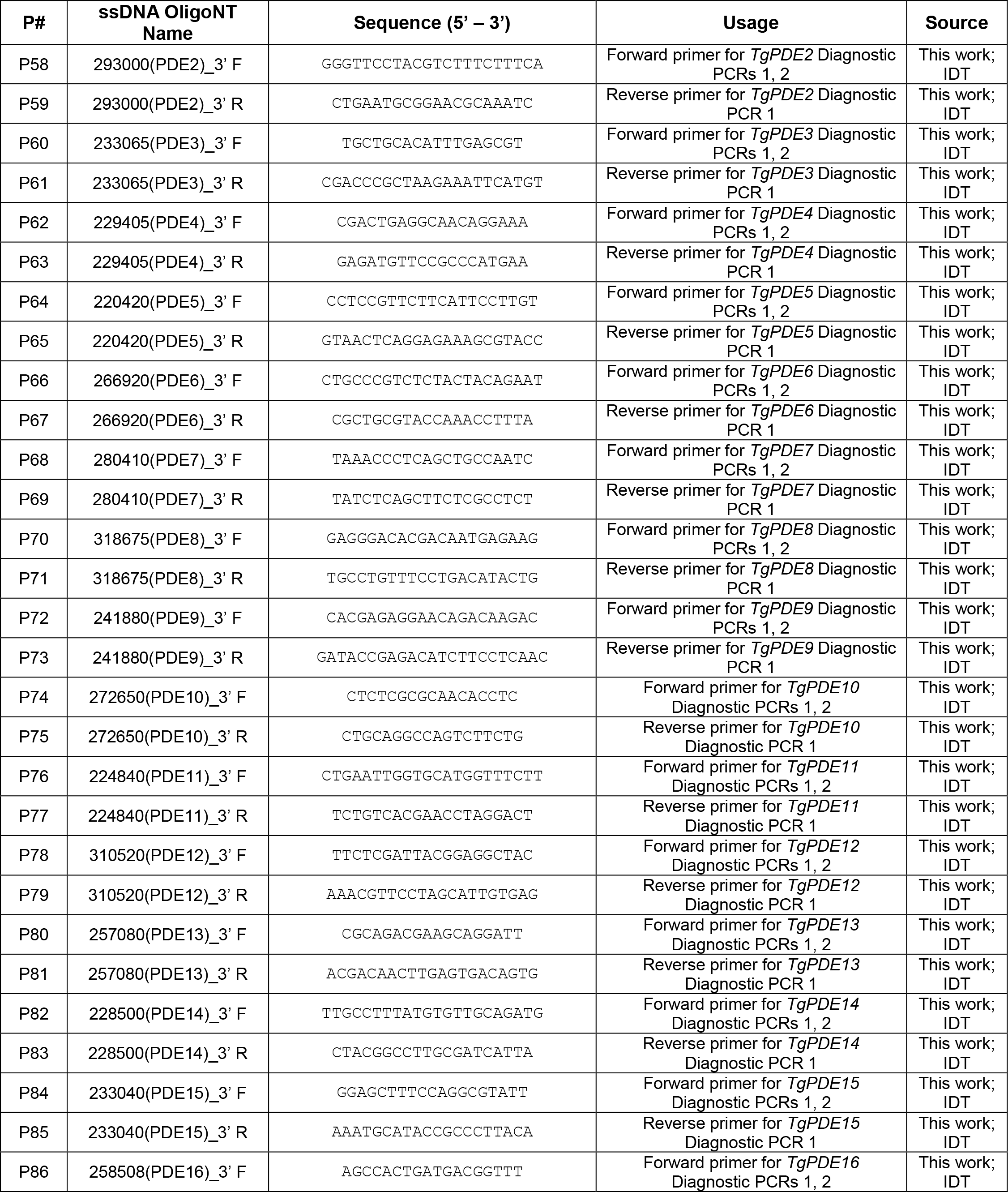
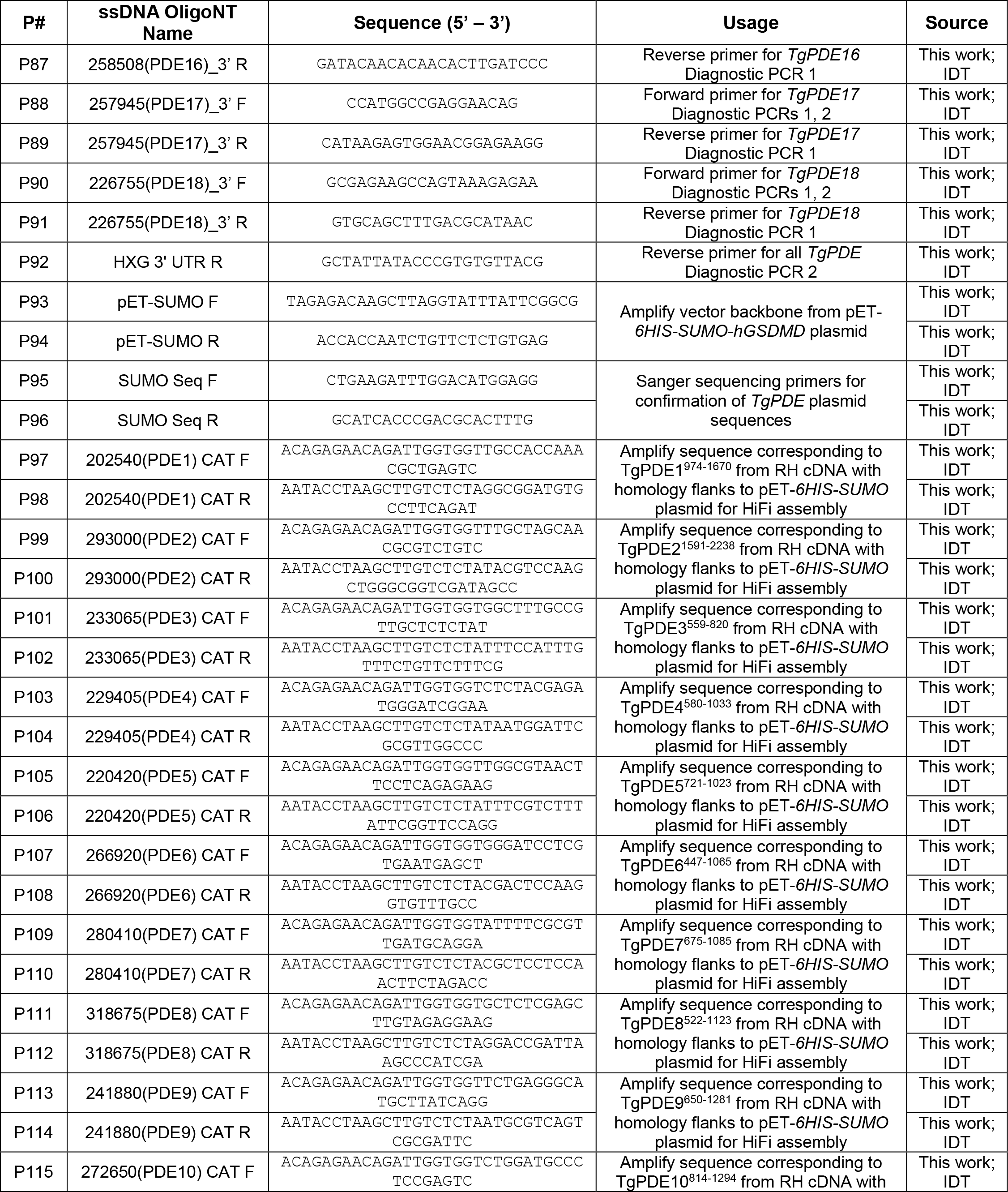
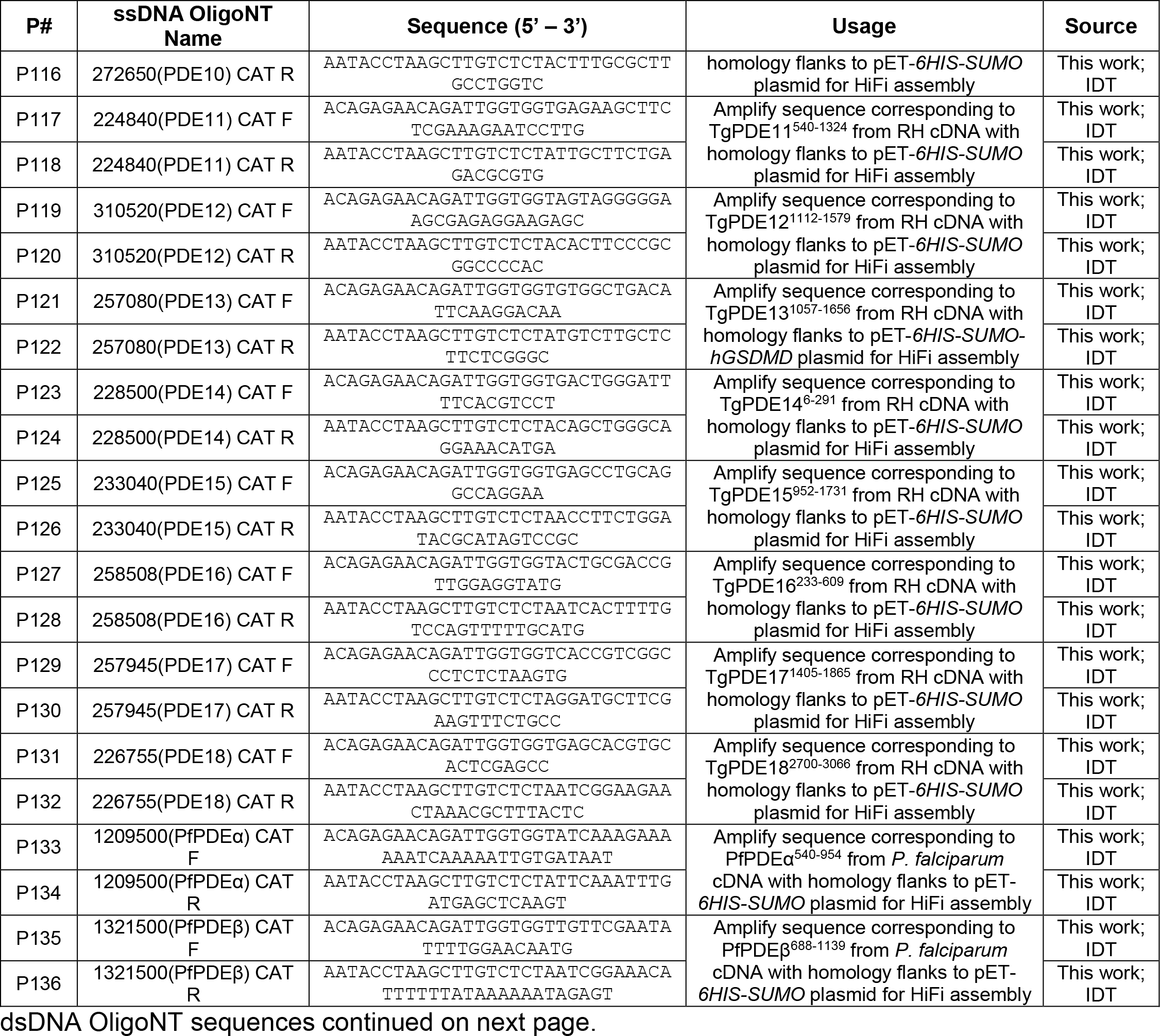
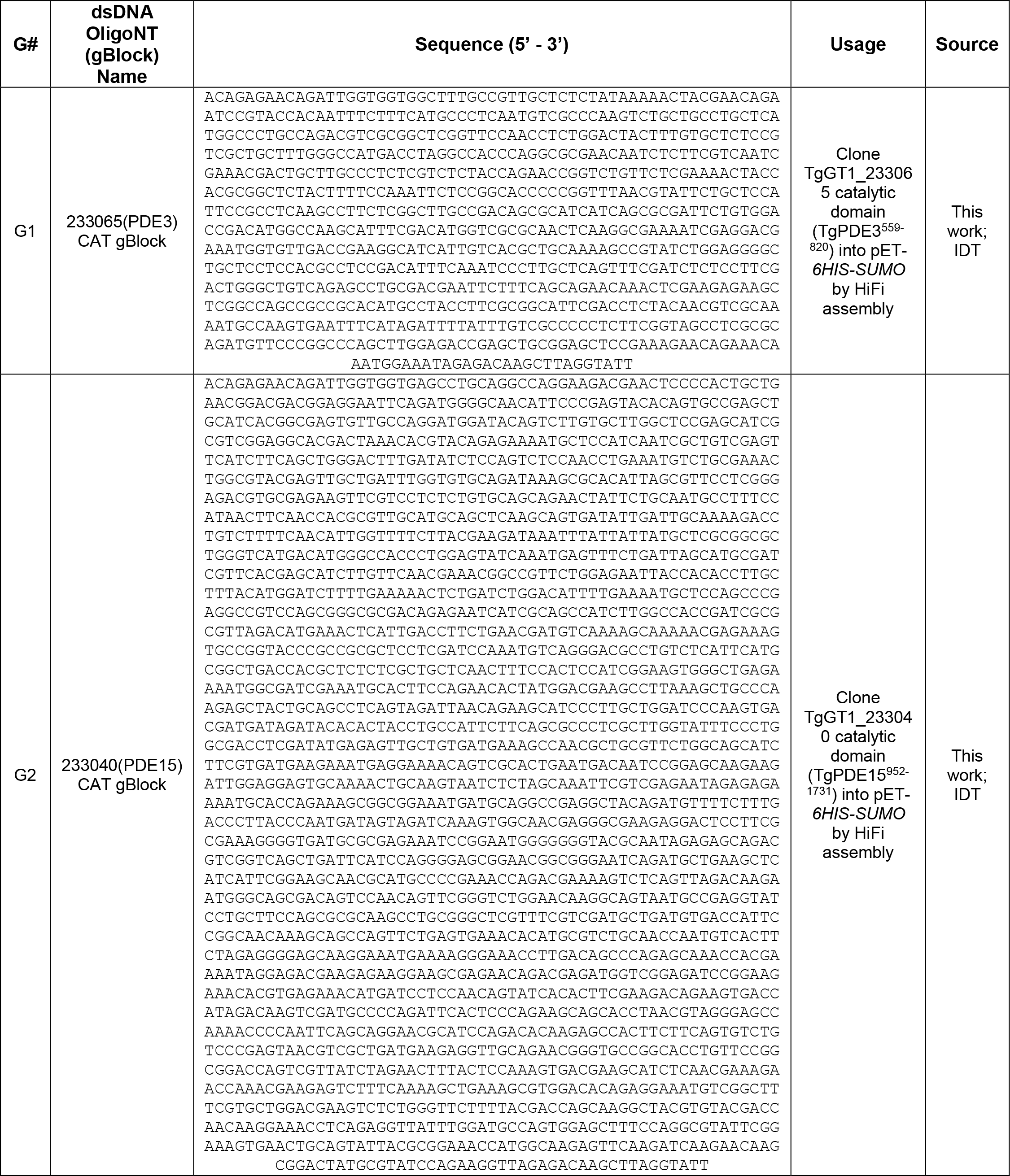
Oligonucleotides used in this study.

## Separate Supplemental Files

Table S4 IP TgPDE1 TgPDE2 LC-MS/MS.xlsx

Table S5 Crosslinked IP TgPDE1 TgPDE2 LC-MS/MS.xlsx

